# The SPARK complex forms the molecular basis of vertebrate fertilization

**DOI:** 10.64898/2026.05.13.724128

**Authors:** Victoria E. Deneke, Johannes P. Suwita, Haoting Wang, Shingo Tonai, Yonggang Lu, Karin Panser, Alexander Schleiffer, Jeremy A. Hollis, Maria Novatchkova, Gerhard Dürnberger, Karel Stejskal, Gabriela Krssakova, Andreas Blaha, Aleq Adrianne R. Andresan, Muriel Mirus, Hana Marvanova, Hsin-Yi Chang, Taichi Noda, Alejandro Burga, Elisabeth Roitinger, Masahito Ikawa, An-drea Pauli

## Abstract

Fertilization requires gamete recognition and membrane fusion, yet the molecular basis of this process in vertebrates remains unknown. Here we identify SPARK (sperm protein assembly and receptor-binding key), a conserved multi-protein complex that integrates all known sperm fertilization factors, including TMEM81–IZUMO1–SPACA6 and DCST1/2, together with two newly identified components, TMDD1 and FAM187A. SPARK subunits are mutually dependent for stability in mature sperm, and disruption of any single component causes male sterility in zebrafish and mice. Incubating zebrafish sperm with soluble egg receptor Bouncer partially rescues fertilization of Bouncer-deficient eggs in a SPARK-dependent manner, consistent with egg receptor binding priming the complex for fusion. Thus, we propose SPARK as a conserved molecular machine that couples gamete recognition to membrane fusion.

## Introduction

The life of every sexually reproducing organism begins with fertilization, the binding and fusion of two gametes. Genetic studies in vertebrates have identified factors on both sperm and egg membranes that are essential for this process (*1–3*). On sperm, these include IZUMO1, SPACA6, TMEM81, DCST1 and DCST2, which are conserved throughout vertebrates and have been shown to be required for fertilization in both fish and mice (*4–12*). In mammals, three additional sperm proteins, TMEM95, FIMP and SOF1, are essential (*8*, *13–17*). On the egg, fertilization requires the GPI-anchored proteins Bouncer (in fish) (*18*) and JUNO (in mammals) (*19*), as well as the tetraspanins CD9 and CD81 (in mammals) (*20–23*).

The sperm protein IZUMO1 directly binds to the egg receptor JUNO, representing the only structurally confirmed interaction between sperm and egg (*24–26*). Consistent with this interaction, sperm-egg binding is impaired in *Izumo1* and *Juno* mutants (*19*, *27*, *28*), while murine sperm lacking any of the other sperm factors are still able to bind to the egg, but fail to fuse (*8*, *9*, *11*, *12*, *14*, *17*). Zebrafish sperm lacking any of the known five sperm factors (IZUMO1, SPACA6, TMEM81, DCST1 and DCST2) are morphologically normal, motile and can approach the micropyle, but fail to bind and fuse to the egg (*10–12*). Thus, in both fish and mammals, loss of any one of these factors results in remarkably similar phenotypes, with sperm failing at the final steps of egg binding and/or fusion.

In some eukaryotes, including plants, algae and protozoans, gamete fusion is mediated by the viral-like fusogen HAP2 (*29–31*). In contrast, no analogous fusogen has been identified in vertebrates. Although IZUMO1 can promote fusion in heterologous systems, it is not sufficient to drive fusion with oocytes (*32*), suggesting that additional factors are required. Thus, a central unanswered question is how sperm and egg factors are organized into a coordinated molecular machinery that mediates egg recognition and membrane fusion in vertebrates.

Recent work has begun to reveal how such coordination may be achieved. TMEM81, IZUMO1 and SPACA6 assemble into a trimeric complex (TIS) that mediates sperm-egg binding by recognizing and adhering to divergent egg receptors – Bouncer in fish and JUNO/CD9 in mammals (*12*, *33*). The receptors both vary between species and also have distinct binding interfaces in fish and mammals: In fish, TIS formation creates the binding interface for Bouncer between the tip of Izumo1 and Spaca6 (*12*). The mammalian TIS complex is predicted to bind to JUNO via a distinct region, the so-called hinge, in IZUMO1 (*12*, *33*), which is consistent with prior structural characterization of the IZUMO1-JUNO interaction (*24–26*). Mammalian TIS is also predicted to bind to the tetraspanins CD9/CD81 on the egg via its subunit IZUMO1 (*33*). While the exact binding mode of TIS to egg receptors differs in fish and mammals, TIS has a conserved function in bridging sperm and egg membranes. In addition, AlphaFold predicts DCST1/2 to form a heterodimer (*12*, *33*), which is in line with published experimental data (*9*, *11*). However, how the remaining sperm fertilization factors contribute to this process remains unknown. In particular, it is unclear whether these proteins act independently or are integrated into a larger molecular assembly (Fig. 1A, B), and how such a system might ultimately drive membrane fusion.

**Fig. 1.**
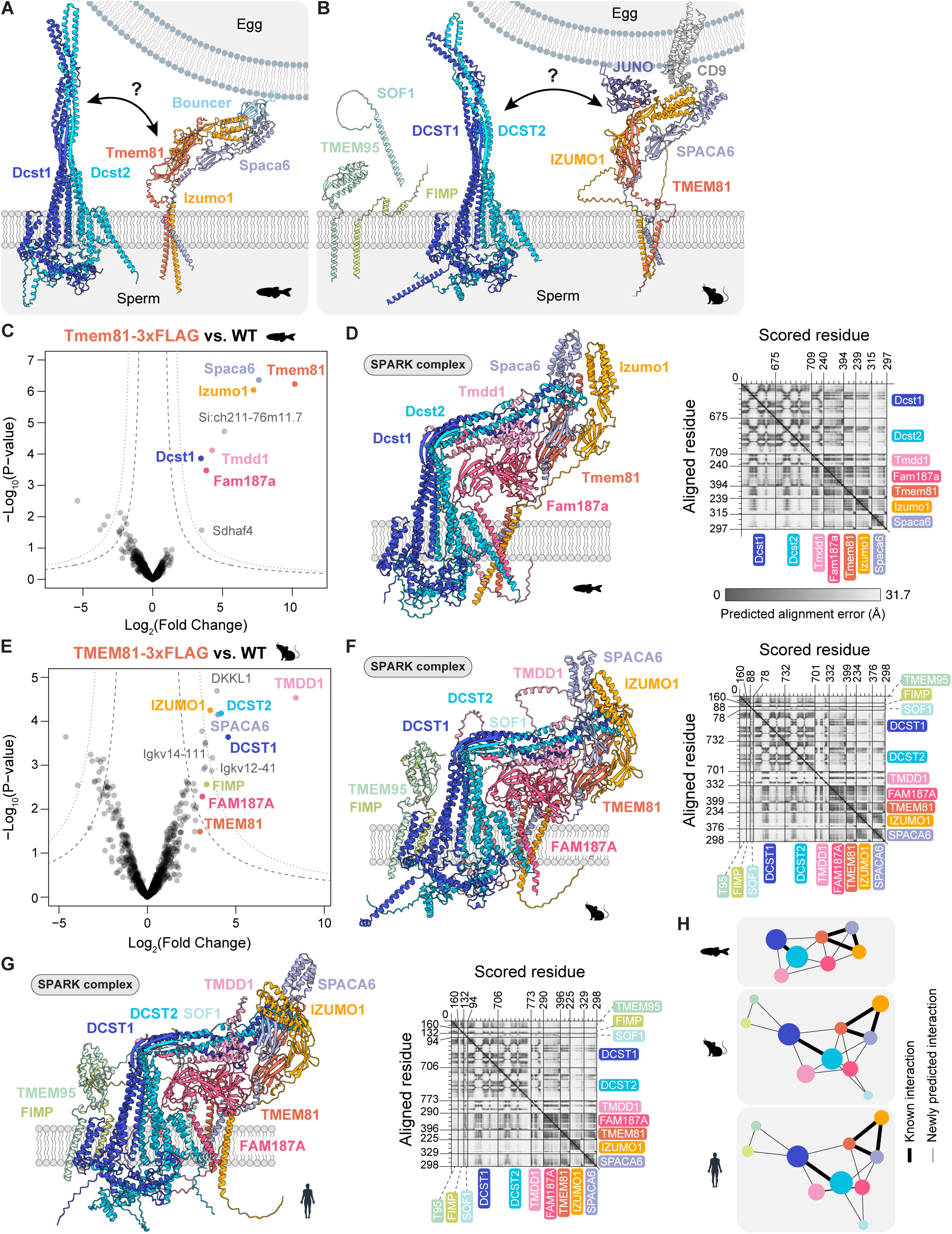
Identification of the SPARK complex. **A**) The zebrafish fertilization synapse. Shown are AlphaFold models of all previously known factors and their interactions on zebrafish sperm and egg. **(B)** The mammalian fertilization synapse. Shown are AlphaFold models of all previously known factors and their interactions on murine sperm and egg. **(C)** Volcano plots of differentially enriched proteins in FLAG co-IP experiments from zebrafish sperm expressing Tmem81-3xFLAG-sfGFP versus wild-type (WT) sperm (based on (*12*)). Fam187a was newly detected by searching against a custom database of ORFs based on zebrafish testis RNA-seq. False discovery rate (FDR) thresholds are indicated as dotted (1% FDR) and dashed lines (5% FDR). Detected SPARK complex members are highlighted in color. **(D)** Zebrafish SPARK complex on the sperm membrane. Left: predicted model; right: predicted alignment error (PAE) plot of the predicted structure. Interface Predicted Template Modeling (iPTM) score: 0.57. **(E)** Volcano plots of differentially enriched proteins in FLAG co-IP experiments from mouse sperm expressing TMEM81-3xFLAG versus wild-type (WT) sperm. False discovery rate (FDR) thresholds are indicated as dotted (1% FDR) and dashed lines (5% FDR). Detected SPARK complex members are highlighted in color. **(F)** Mouse SPARK complex on the sperm membrane. Left: predicted model; right: predicted alignment error (PAE) plot of the predicted structure. iPTM: 0.53. **(G)** Human SPARK complex on the sperm membrane. Left: predicted model; right: predicted alignment error (PAE) plot of the predicted structure. iPTM: 0.51. **(H)** Comparison of the interaction network of the zebrafish, mouse and human SPARK complexes. Nodes represent individual subunits (colors as indicated in previous panels). Bold lines highlight the two known subcomplexes DCST1/2 and TMEM81-IZUMO1-SPACA6 within the SPARK complex. All other interactions (thin lines) were newly predicted.

Here, we show that all known vertebrate sperm fertilization factors together with two newly identified proteins, TMDD1 and FAM187A, form the Sperm Protein Assembly and Receptor-binding Key (SPARK) complex. Egg receptor binding is required for priming sperm-egg fusion, suggesting a mechanism by which the SPARK complex couples sperm-egg recognition and membrane fusion.

## Results

### Identification of the SPARK complex

To determine whether vertebrate sperm fertilization factors form a larger molecular assembly, we reanalyzed a previously published Tmem81 co-immunoprecipitation mass spectrometry (co-IP/MS) dataset from zebrafish sperm (*12*) (Table S1). In addition to the TIS complex, Dcst1 was significantly enriched (Fig. 1C), providing initial evidence for an interaction between the two sperm complexes Dcst1/2 and Tmem81-Izumo1-Spaca6 (TIS) (Fig. 1A, B). In addition to Dcst1, three other proteins with no prior links to fertilization were also significantly enriched in the Tmem81 co-IP/MS experiment (Fig. 1C; Tmdd1, Si:ch211-76m11.7, Sdhaf4). To test whether any of these enriched factors might directly interact with TIS and/or Dcst1/2, we used AlphaFold Multimer to predict pairwise interactions (*34–37*). Tmdd1 (Transmembrane and death domain protein 1) stood out as the highest-scoring interactor for both Dcst1/2 and TIS (fig. S1A, B). When predicting all six proteins together, Tmdd1 was predicted to form a complex with Dcst1/2 (fig. S1C, S1D). However, the interaction between this Tmdd1-Dcst1/2 trimer and TIS was less confidently predicted (fig. S1C). Given that both Dcst1 and Tmdd1 were significantly enriched in the Tmem81-3xFLAG IP, we reasoned that an additional factor might be required to bridge these two trimers.

Motivated by our previous discovery of Bouncer as an unannotated protein-coding gene (*18*), we asked whether our co-IP/MS dataset might contain peptides derived from unannotated regions of the zebrafish genome. To address this, we assembled transcripts from zebrafish testis RNA-seq data (*18*) using both reference-guided and *de novo* approaches. We then searched for peptides identified in the Tmem81-3xFLAG IP against predicted open reading frames (ORFs) derived from these transcripts (Table S1), revealing a conserved ORF not currently annotated in the zebrafish genome. This ORF encodes a homologue of mammalian FAM187A, a single-pass type I transmembrane protein with an Ig-like V-type ectodomain. Quantification of corresponding peptides showed that Fam187a is significantly enriched in the Tmem81 IP (Fig. 1C). Similarly, reciprocal co-IP/MS experiments using Spaca6-3xV5 as an independent bait also enriched Tmdd1 and Fam187a, supporting their identification as *bona fide* TIS interactors in sperm (fig. S1E, Table S1). Notably, inclusion of Fam187a in addition to TIS, Dcst1/2 and Tmdd1 in AlphaFold predictions revealed a single large multimeric protein complex (Fig. 1D, Supplementary Movie 1). Because this predicted structure integrates all known fertilization factors along with two newly identified components, we refer to this assembly as the Sperm Protein Assembly and Receptor-binding Key (SPARK) complex.

We detected transcripts encoding the two newly identified factors, Tmdd1 and Fam187a, in RNA-seq data from testes of both zebrafish and mice (fig. S1F, G), suggesting that the SPARK complex may be conserved in mammals. To test this, we performed an analogous co-IP/MS experiment using transgenic mouse sperm expressing TMEM81-3xFLAG. All SPARK complex members identified in zebrafish - TIS, DCST1/2 and the two newly identified factors TMDD1 and FAM187A - were enriched, along with FIMP (Fig. 1E, Table S1). AlphaFold predicted that these eight proteins, together with the additional mammalian fertilization factors TMEM95 and SOF1, assemble into a ten-subunit SPARK complex (Fig. 1F, Supplementary Movie 1). Multiple lines of evidence indicate that this complex is also conserved in humans: transcripts for *Tmdd1*, *Fam187a* and all other SPARK components are expressed in the human testis (*38*). Moreover, AlphaFold predicts a structurally analogous complex formed by the human orthologs (Fig. 1G, Supplementary Movie 1), which is consistent with recent proteome-wide predictions of protein-protein interactions that found human FAM187A as an interactor of both TMEM81 and DCST2 (*39–41*). Comparison of the zebrafish, mouse and human assemblies revealed that the seven-subunit SPARK complex in fish constitutes a conserved core that is also present in mammals (Fig. 1H).

### SPARK complex protein levels are co-dependent in vertebrate sperm

To further test whether the SPARK subunits form a protein complex *in vivo*, we examined whether SPARK protein levels are co-depleted upon loss of any single component. Given the limited availability of antibodies for SPARK members and their relatively low abundance, we employed Parallel Reaction Monitoring Mass Spectrometry (PRM-MS). This enabled detection of all SPARK complex members in wild-type zebrafish sperm (Fig. 2A). We then quantified protein levels of each SPARK member across all previously published SPARK member mutants (*10–12*) and newly generated tmdd1 and fam187a mutants (Fig. 2A; fig. S2A, B; Table S1). Importantly, all SPARK proteins were depleted in each single-knockout (KO) sperm sample (Fig. 2A), suggesting that the stability of each component depends on the formation of the complete complex in mature zebrafish sperm.

**Fig. 2.**
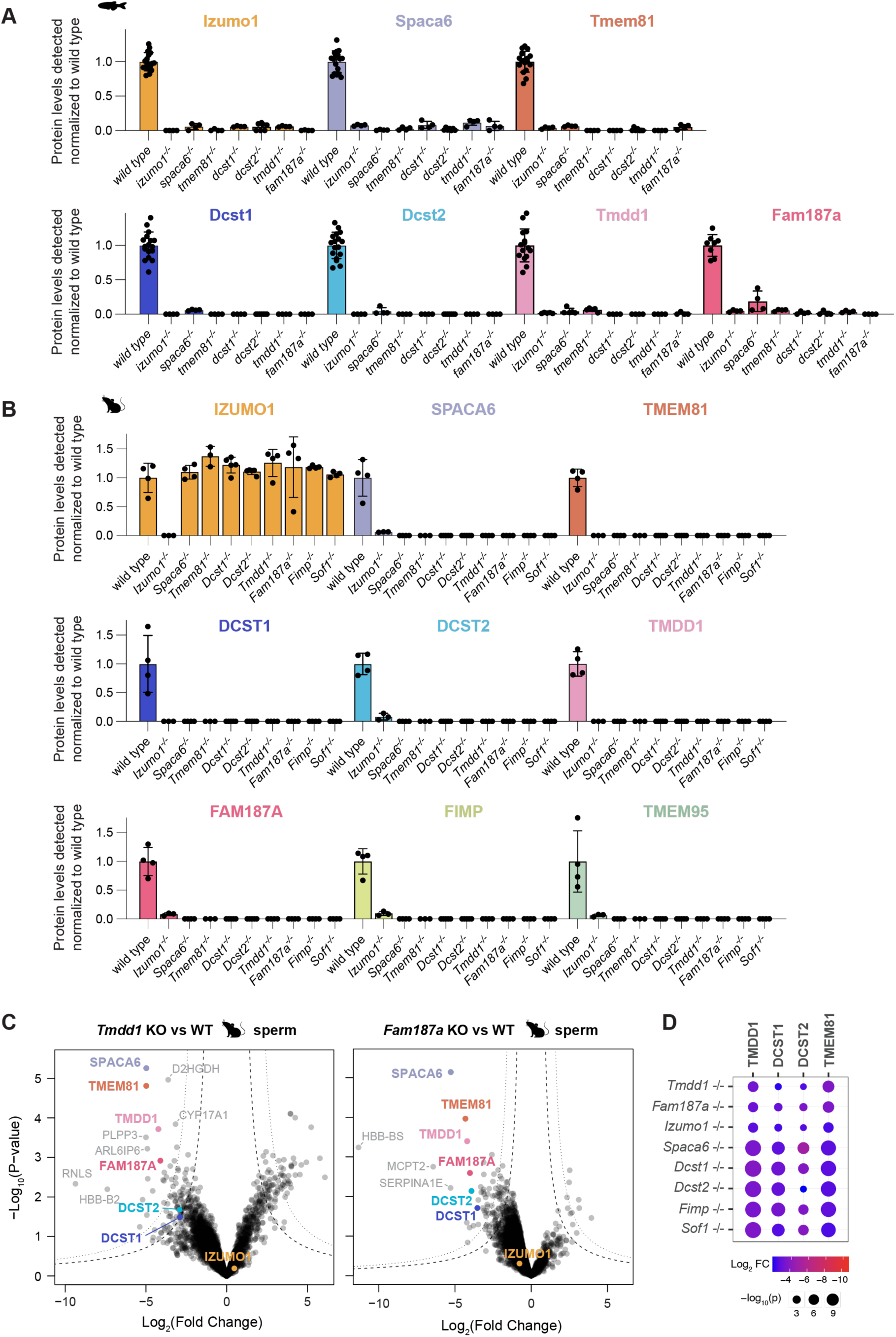
SPARK complex members are co-dependent and co-depleted in vertebrate sperm. **(A)** Relative protein levels of SPARK complex members in wild-type and mutant zebrafish sperm by PRM-MS normalized to six constant proteins. All comparisons statistically significant, see Table S1 for calculated p-values (permutation test). **(B)** Relative protein levels of SPARK complex members in wild-type and mutant mouse sperm by PRM-MS normalized to four constant proteins. SOF1 could not be detected by PRM–MS due to lack of suitable peptides. All comparisons statistically significant except IZUMO1, see Table S1 for calculated p-values (permutation test). **(C)** Volcano plots of differentially regulated proteins in wild-type (WT) versus *Tmdd1* KO (left) and *Fam187a ^Δ2^* KO (right) sperm samples. Detected SPARK complex members are highlighted in color. False discovery rate (FDR) thresh-olds are indicated as dotted (1% FDR) and dashed lines (5% FDR). **(D)** Bubble plot of significantly depleted proteins (Log2 Fold Change < -2, *p*-value < 0.05) in mouse sperm samples of all measured genotypes. Color gradient indicates the magnitude of depletion (Log2 Fold Change), bubble size corre-sponds to the statistical significance (-log10(*p*)). Proteins and genotypes were organized using hierarchical clustering (Euclidean distance, complete linkage) based on their Log2 Fold Change profiles.

To determine whether the protein levels of mammalian SPARK components are also co-dependent, we followed a similar approach for all previously published mutants (*4*, *7–9*, *11*, *12*, *14*) and newly generated *Tmdd1* and *Fam187a* mutant backgrounds (fig. S2C, D; Table S1). Consistent with the co-dependence of SPARK members observed in zebrafish sperm, all SPARK complex proteins detectable in wild-type murine sperm were depleted in single-knockout (KO) mature sperm, with the exception of IZUMO1 (Fig. 2B). The stability of murine IZUMO1 has been noted previously in some mutant backgrounds (*7*, *9*, *11*, *14*), and our analysis extends these findings by showing that IZUMO1 is also not depleted in *Tmem81-/-, Tmdd1-/-* and *Fam187a-/-* mutants (Fig. 2B). Additionally, in line with prior reports in a subset of mutants (*9*, *42*), we find murine SPACA6 depletion in sperm across all mutant backgrounds. In contrast to the observed co-depletion of protein levels in SPARK mutants, SPARK protein levels were not reduced in a SPARK-unrelated *Cd46* mutant background (fig. S3A) (*43*), indicating that this co-dependence is specific to SPARK complex members rather than a general consequence of disrupting acrosomal membrane proteins. SPARK proteins were present in single mutant testis samples (fig. S3B, C), suggesting that they were produced during early spermatid stages, independent of the presence of other SPARK members, but were depleted in mature sperm (Fig. 2B).

To globally assess whether any proteins other than SPARK members may be co-depleted in SPARK mutants, we performed shotgun mass spectrometry on wild-type and SPARK member mutant mouse sperm (Fig. 2C; fig. S3D, Table S1). Although other proteins showed partial co-depletion across genotypes (fig. S3E), the only proteins consistently detected by shotgun MS and depleted across all assessed mutant genotypes were the four SPARK components TMDD1, TMEM81, DCST1 and DCST2 (Fig. 2D, fig. S3E). Together, these data support a model in which the SPARK complex comprises seven core members in fish and ten complex members in mammals.

### TMDD1 and FAM187A are essential for fertilization

Having shown that the two newly identified factors TMDD1 and FAM187A are core members of the SPARK complex in both fish and mammals, we set out to determine if these factors are essential for fertilization. Analogous to all other SPARK subunit mutants in fish (*10–12*) and mice (*4*, *7–9*, *11*, *12*, *14*), males lacking *Tmdd1* or *Fam187a* were sterile in both organisms (Fig. 3A, C-D). In zebrafish, mutant sperm were motile and could reach the egg (fig. S4A, B), but failed to stably adhere to the egg plasma membrane (Fig. 3B; Supplementary Movie 2), in line with the phenotypes reported for other SPARK mutants in fish (*10–12*). In mice, *Tmdd1* and *Fam187a* KO sperm exhibited normal progressive motility, underwent the acrosome reaction (fig. S4C, Fig. 3E) and were able to bind the oolemma (Fig. 3E), but failed to fuse with the egg (Fig. 3F), in agreement with previously described SPARK mutant phenotypes (*4*, *7–9*, *11*, *12*, *14*). Overall, our results establish TMDD1 and FAM187A as essential factors for vertebrate fertilization.

**Fig. 3.**
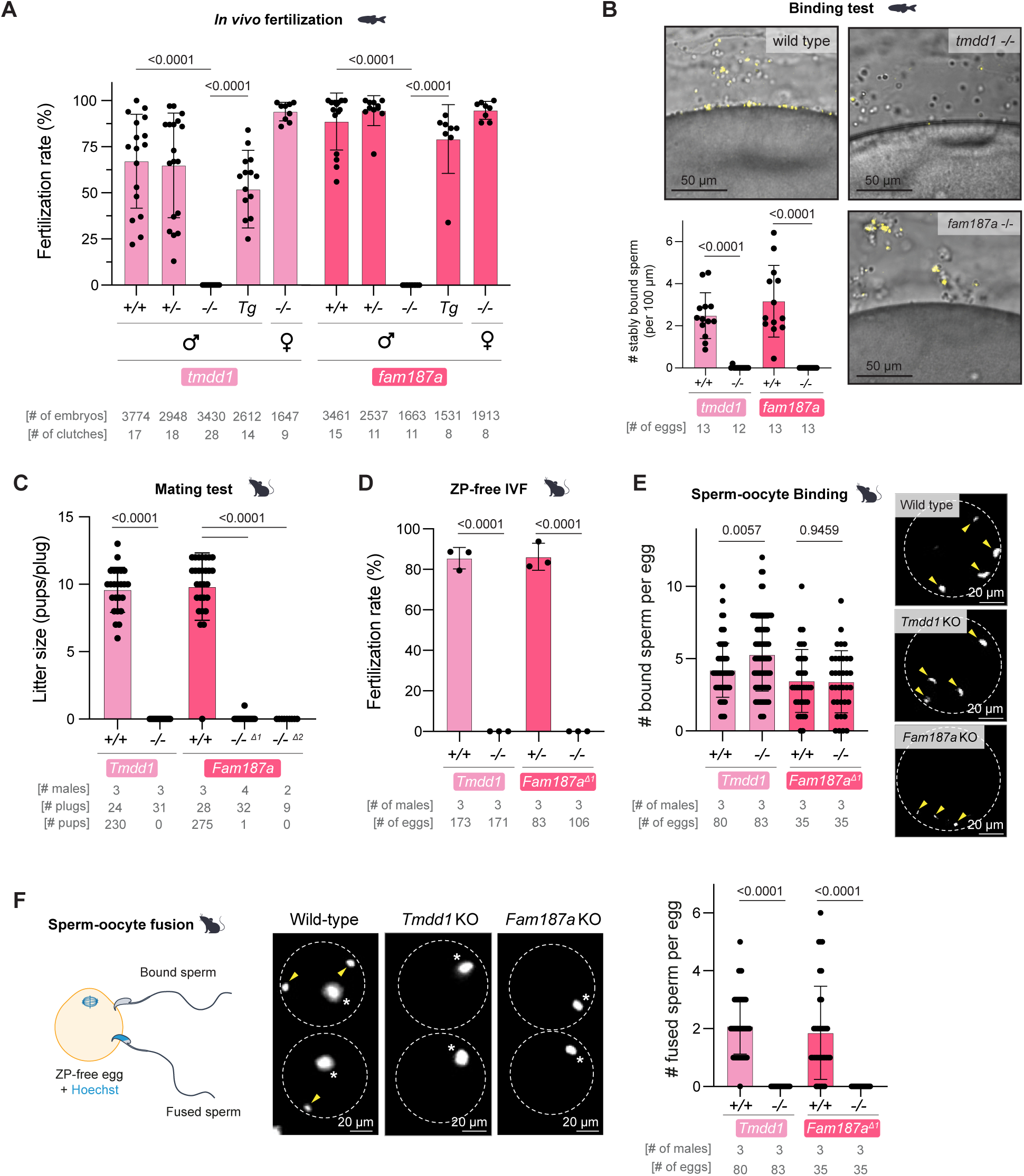
TMDD1 and FAM187A are essential for male fertility in zebrafish and mice. **(A)** *In vivo* fertilization rates in zebrafish. Males or females of the indicated genotypes (+/+, wildtype; +/-, heterozygous; -/-, KO) were crossed with wild-type zebrafish. *Tg*: *tmdd1* KO (−/−) fish expressing transgenic Tmdd1-3xFLAG-mScarlet3 and *fam187a KO* (−/−) fish expressing transgenic Fam187a-3xFLAG-sfGFP, respectively. **(B)** Sperm-egg binding assay in zebrafish. Sperm labeled with MitoTracker (yellow) was incubated with activated, dechorionated wild-type eggs. Representative images of wild-type, *tmdd1* KO, and *fam187a* KO sperm binding to the surface of the egg 1 min after sperm addition. Bottom left: quantification of stably bound sperm per 100 μm over a period of 2 min. **(C)** Average litter size after 8-week pairings of mutant male mice of the indicated genotypes (+/+, wildtype; -/-, KO) with wild-type female mice. **(D)** *In vitro* fertilization rates using wild-type, *Tmdd1* KO or *Fam187a* KO murine sperm and ZP-free wild-type eggs. **(E)** Sperm-oocyte binding assay in mice. Left: quantification of total number of acrosome-reacted sperm bound to the oolemma. Right: representative images of wild-type, *Tmdd1* KO and *Fam187a* KO sperm stained for IZUMO1 (white). Yellow arrowheads: acrosome-reacted sperm. **(F)** Sperm-oocyte fusion assay in mice by Hoechst transfer into sperm from pre-loaded eggs. Left: scheme of the assay. Middle: representative images of the sperm-oocyte fusion assay in mice; yellow arrowheads: fused sperm; asterisks: egg meiosis II chromosomes. Right: quantification of number of fused sperm per egg. Bar graphs represent mean ± SD. *P*-values are calculated by Kruskal-Wallis test with Dunn’s multiple comparisons test (A, C (*Fam187a* panel)), or Mann-Whitney test (B, C (*Tmdd1* panel), D-F).

### Evolutionary conservation and origin of the SPARK complex

To assess whether the newly identified factors are present beyond vertebrates, we first examined their taxonomic distribution and found that TMDD1 and FAM187A are conserved across diverse bilaterian lineages (Fig. 4).

**Fig. 4.**
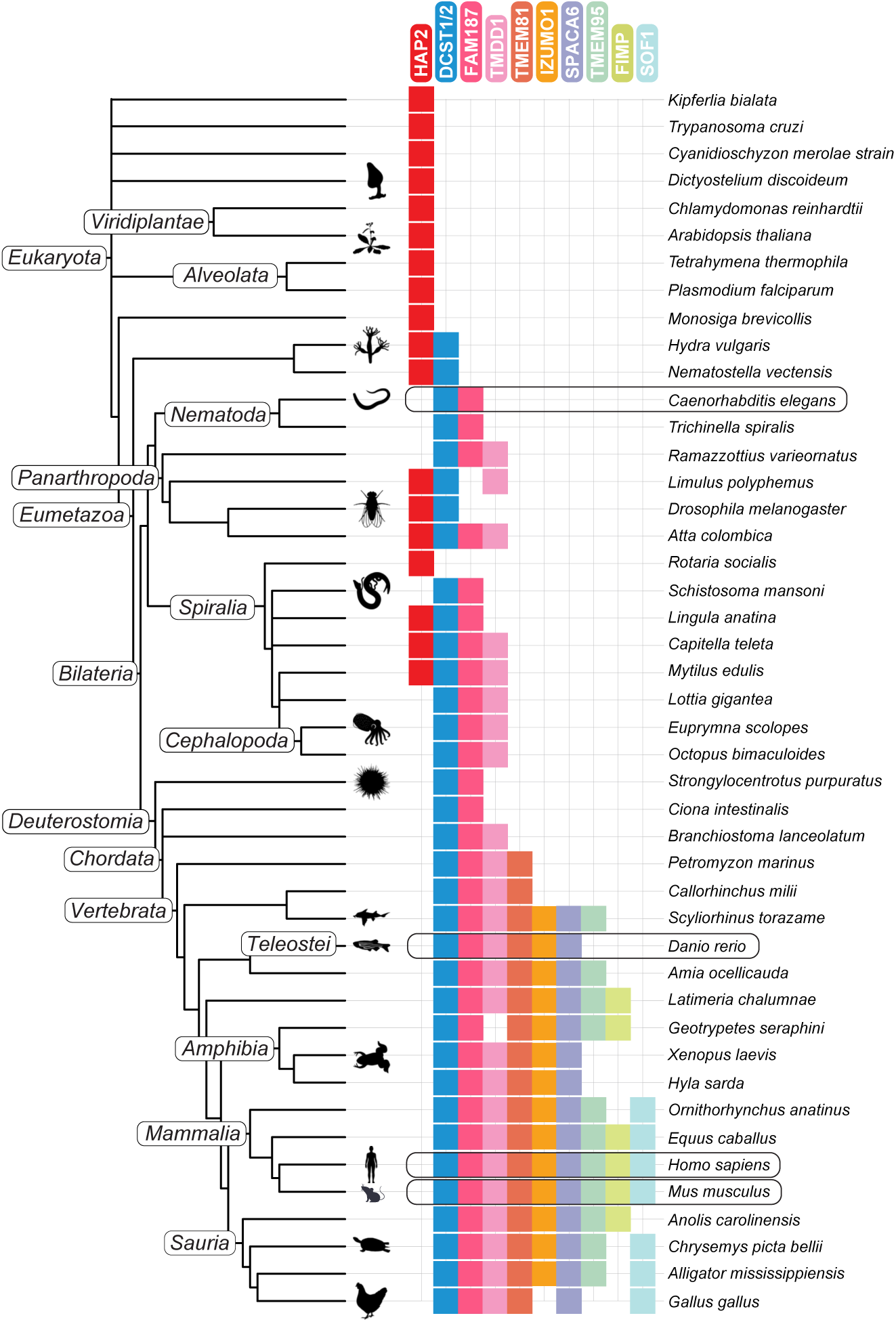
Evolutionary conservation and origin of the SPARK complex. The taxonomic tree follows the NCBI taxonomy. Filled boxes indicate significant blast hits in NCBI nr or UniProt ref-erence proteomes using a family-specific reference sequence set. Two paralogous families, DCST1 and DCST2, and FAM187A and FAM187B, cannot be subclassified unambiguously in all lineages and are combined. The *Danio rerio* Fam187a protein sequence has been submitted to the NCBI gene database as part of this study. Silhouettes were reproduced from PhyloPic (https://www.phylopic.org/).

To test whether this conservation extends to function, we focused on FAM187A (fig. S5A), which has a clear ortholog in *C. elegans*, *fam187/spe-40* (Y37E11AR.7) (Fig. 4, fig. S5B), and is expressed in spermatids and mature sperm (*44*). We generated a null mutant (fig. S5B) and assessed fertility by measuring brood size. While heterozygous *fam187/spe-40* mutants produced progeny comparable to wild type, homozygous mutants were sterile (fig. S5C). This sterility was rescued by mating with wild-type males, indicating that the defect is sperm-specific (fig. S5C). These results are consistent with a recent independent study (*45*) and demonstrate that FAM187 has a conserved and essential role in sperm-mediated fertilization in *C. elegans*.

In addition to TMDD1 and FAM187A, DCST1 and DCST2 stand out as even more broadly conserved SPARK complex members (*12*, *46*). Both proteins are not only phylogenetically conserved, but also functionally, as they are required for fertilization in the invertebrates *C. elegans* and *D. melanogaster* (*47–51*). In contrast, the remaining SPARK components are restricted to vertebrate lineages: IZUMO1, SPACA6, and TMEM81 are broadly conserved across vertebrates, whereas TMEM95, FIMP, and SOF1 show a more limited and scattered distribution (Fig. 4).

Taken together, these findings suggest that TMDD1 and FAM187A, along with DCST1 and DCST2, represent some of the most ancient and evolutionarily conserved components of the sperm fertilization machinery. We therefore propose that the SPARK complex originated from a DCST1/2 core, subsequently expanded to include FAM187A and TMDD1, and later incorporated additional vertebrate-specific components.

Fertilization components are predicted to be under intense selective pressure from gamete competition at sperm-egg interfaces. Given this, as well as the presence of vertebrate-specific fertilization factors, we expected SPARK components to display signatures of positive, or diversifying, selection. PAML analyses identified positive selection in 5/10 *Murinae* components tested (fig. S5D; Table S2), with most selection concentrated at the distal tip of SPARK (fig. S5D-G), consistent with this region physically interacting with the egg surface (*12*, *19*, *33*). Extending previous studies on signatures of positive selection within IZUMO1 and JUNO (*52*, *53*), we found reciprocal selection in egg components CD9 and JUNO (fig. S5H-I), further supporting a physical interaction between these factors and lineage-specific selective pressure.

### The SPARK complex is activated by Bouncer binding preceding sperm-egg fusion

Beyond Bilateria, sperm-egg fusion is driven by the viral-like fusogen HAP2/GCS1 in plants, algae and protozoans (*31*, *54–57*). HAP2 is widely conserved in plants and protists, and also present in some metazoans, sharing limited overlap with DCST1/2 (Fig. 4; fig. S5J) (*46*).

Viral and viral-like fusogens, such as HAP2, often mediate membrane fusion by binding to receptor proteins on the target membrane. This interaction can induce conformational rearrangements in the fusogen that precede membrane fusion (*58*, *59*). We hypothesized that - in analogy to viral and viral-like fusogens - egg receptor binding (Bouncer in fish and JUNO/CD9 in mammals) might similarly activate the SPARK complex for fusion (Fig. 5A). Indirect support for this hypothesis comes from *in vitro* studies demonstrating that IZUMO1 undergoes conformational changes upon binding to JUNO (*24*, *60*). In fish, Bouncer is predicted to interact with the SPARK complex through the 4-helix bundle (4HB) domains of both Spaca6 and Izumo1 (*12*). AlphaFold models of the Bouncer-bound state of the SPARK complex suggest that Bouncer binding brings the 4HB domains of Spaca6 and Izumo1 into closer proximity (Fig. 5B), accompanied by conformational rearrangements in the SPARK complex (Supplementary Movie 3). To assess whether Bouncer binding alters the interactions within the SPARK complex, we incubated zebrafish sperm expressing Spaca6-3xV5 either with recombinant Bir2-Bouncer or with the Bir2 solubility tag alone (negative control). Co-IP/MS using Spaca6-3xV5 as bait revealed that most SPARK members were equally enriched in both conditions, but Izumo1 was significantly more enriched in the presence of Bouncer compared to Bir2, suggesting that receptor binding indeed enhances the interaction between Spaca6 and Izumo1 (fig. S6A, Table S1).

**Fig. 5.**
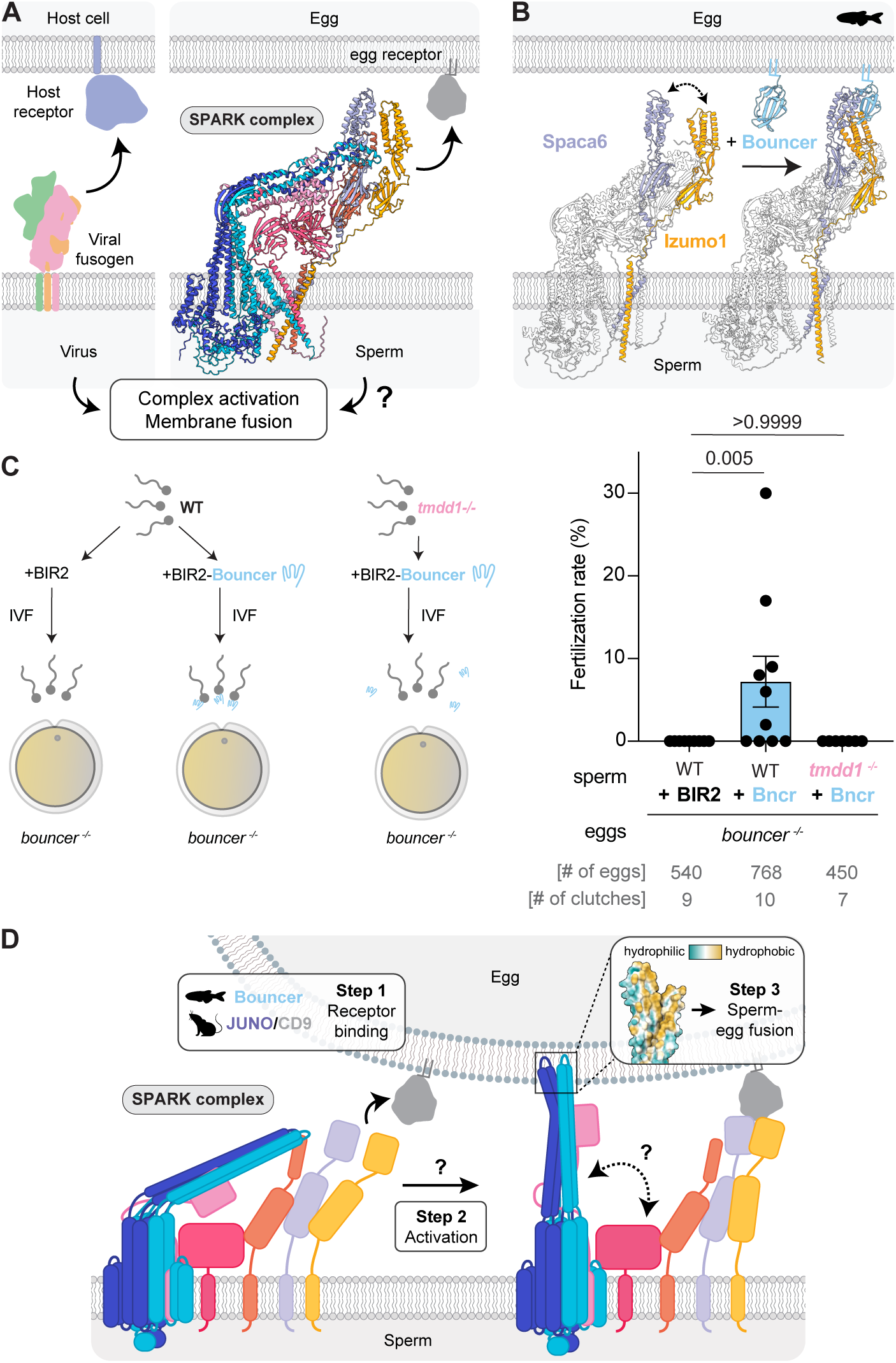
The SPARK complex is activated by egg receptor binding preceding sperm-egg fusion. **(A)** Viral fusogens recognize surface receptors on the host cell membrane, followed by activation of the fusogen complex and membrane fusion. Similarly, the SPARK complex binds to an egg receptor. In parallel to viral fusogens, following receptor binding the two membranes fuse. **(B)** Structural predictions of the zebrafish SPARK complex with and without the zebrafish egg receptor Bouncer. Bouncer binding is predicted to lead to a conformational change in the SPARK complex, bringing Spaca6 and Izumo1 in closer proximity and allowing Bouncer binding to the 4-helix bundle domains of both Spaca6 and Izumo1. **(C)** Soluble recombinant Bouncer is sufficient to activate SPARK-containing zebrafish sperm to enable sperm-egg fusion in the absence of membrane-bound Bouncer. Left: experimental setup. Zebrafish sperm (wild type (WT) or *tmdd1 ^-/-^*) was pre-incubated with BIR2 (solubility tag) or BIR2-Bouncer and added to *bouncer ^-/-^* eggs. Right: *In vitro* fertilization rates. Bar graphs represent mean ± SD. *P*-values are calculated by Kruskal-Wallis test with Dunn’s multi-ple comparisons test. **(D)** Model for SPARK-mediated sperm-egg binding followed by membrane fusion. In Step 1, the SPARK complex binds to the egg receptor (Bouncer in fish and JUNO/CD9 in mammals). In Step 2, the SPARK complex is activated by un-dergoing a conformational change, releasing the hydrophobic tip of DCST1/2 and leading to its insertion into the egg membrane. This destabilizes the egg membrane and ultimately leads to membrane fusion (Step 3).

We next asked whether receptor-binding to the SPARK complex is able to activate sperm for fusion. To test this, we incubated wild-type sperm with soluble Bir2-Bouncer or Bir2 alone and assessed the sperm’s ability to fertilize *bouncer^-/-^* eggs. Sperm incubated with Bir2 alone failed to fertilize *bouncer*^-/-^ eggs (Fig. 5C). Notably, sperm pre-incubated with soluble BIR2-Bouncer fertilized *bouncer* ^-/-^ eggs at low but statistically significant rates (median ∼7%, range 0-30%; p=0.005, Fig. 5C). This effect was dependent on the SPARK complex, as *tmdd1* KO sperm pre-incubated with Bir2-Bouncer were unable to fertilize *bouncer* ^-/-^ eggs (Fig. 5C). Together, these results suggest that binding of the egg receptor Bouncer can prime the zebrafish SPARK complex for fusion. This activation can occur independently of membrane anchoring, despite Bouncer being normally GPI-anchored on the egg surface (*18*). Consistent with this, pre-incubation of mouse sperm with CD9-containing vesicles shed from wild-type oocytes was previously reported to rescue fertilization of *Cd9* KO oocytes (*61*), although this observation has not been consistently reproduced (*62*, *63*). We therefore propose that egg receptors serve a dual role in fertilization: they first tether sperm via SPARK binding and subsequently activate the complex at the plasma membrane, where proximity to lipids may facilitate full conformational rearrangements leading to fusion. This model is consistent with recent ultrastructural studies of fertilization mutants (*28*) and mechanistic characterization of the fertilization-blocking antibody OBF13 (*64*), suggesting that IZUMO1–JUNO primarily mediate gamete recognition, whereas other SPARK components may be required for fusion initiation.

If receptor binding represents the initial activation step, how is the SPARK complex subsequently rearranged to initiate membrane fusion? We predicted the DCST1/2 dimer from different species which consistently showed that the stalk-like DCST1/2 ectodomain exposes hydrophobic residues at its tip (fig. S6B), a feature reminiscent of the hydrophobic loops used by viral(-like) fusogens, including HAP2, to initiate membrane insertion and destabilization (*29–31*, *58*, *59*). Positive selection analysis revealed that this region also contains a high concentration of residues under diversifying selection, much like viral fusogen proteins (fig. S5D, G). Within the SPARK complex, however, DCST1/2 are predicted to adopt a bent conformation in which this hydrophobic tip is shielded through interaction with the N-terminus of TMEM81 (fig. S6C, D; Supplementary Movie 4), similar to the proposed shielding function of the A16/G9 interaction in the vaccinia virus entry fusion complex (*65*).

We therefore propose that egg receptor binding (in proximity to the egg membrane) induces a conformational rearrangement in the SPARK complex that leads DCST1/2 to adopt an extended conformation, exposing the hydrophobic tip for interaction with the egg plasma membrane. Analogous to the pre-hairpin intermediate of viral fusogens (*58*, *59*), insertion of these hydrophobic residues could destabilize the egg membrane and promote fusion (Fig. 5D). Together, our findings implicate the SPARK complex as the molecular machinery on sperm that couples sperm-egg binding and fusion in vertebrates.

## Discussion

Here, we identify the SPARK complex, a conserved multi-subunit assembly that contains all known vertebrate sperm fertilization factors and the two newly identified factors, TMDD1 and FAM187A (Fig. 1C-F), which collectively mediates sperm-egg binding and fusion (Fig. 5D). The vertebrate SPARK complex comprises seven core factors and is predicted to include three additional factors in mammals (Fig. 1D, F-H). While our AlphaFold models and mutant proteomic data support a ten-factor assembly in mammals, SOF1 and TMEM95 were not detected in our co-IP–MS experiments (Fig. 1E). Nonetheless, TMEM95 was co-depleted in all SPARK member mutants, which is in line with a reported interaction between FIMP and TMEM95 (*66*), further supporting that TMEM95 is part of this assembly *in vivo*. Similarly, *Sof1* mutants also show co-depletion of all SPARK members (Fig. 2B); however, whether SOF1 is a stable component of the mammalian SPARK complex in mature sperm remains unresolved. More broadly, although AlphaFold strongly predicts a human SPARK complex, its existence in human sperm remains to be experimentally validated.

Analysis of SPARK protein co-dependence revealed a largely conserved pattern with one notable exception: in zebrafish, all components are mutually dependent, whereas in mice IZUMO1 is uniquely retained in the absence of other factors (Fig. 2A, B). This may reflect its ability to homodimerize (*67*), potentially allowing a fraction of IZUMO1 to remain stable outside the complex. Whether this difference reflects a broader feature of mammalian IZUMO1 biology or an additional functional role remains to be tested in the future.

Our study uncovers two previously uncharacterized factors essential for fertilization - TMDD1 and FAM187A. TMDD1 is a single-pass type I membrane protein with an extracellular death domain (fig. S6E), a unique feature among known death domain–containing proteins in vertebrates (fig. S6F, G). Death domains are known to mediate oligomerization and higher-order assembly (*68*, *69*), consistent with TMDD1 being a central member of the SPARK complex. FAM187A contains an Ig-like V-type domain and has a conserved role in fertilization also in invertebrates (fig. S5B, C) (*45*). Together, these features suggest that FAM187A may represent an early adhesion module within the fertilization machinery.

We speculate that during fertilization, the SPARK complex needs to undergo an activation step which leads to sperm-egg fusion. An important insight from our soluble recombinant experiments is that egg receptor binding serves as more than a docking factor for the sperm complex. Specifically, we find that soluble Bouncer is sufficient to enable zebrafish sperm to fuse with Bouncer-deficient eggs (Fig. 5C). However, this contrasts with earlier observations that recombinant Bouncer reduces fertilization rates in wild-type eggs (*12*), indicating that its effect is strongly context dependent.

The nature of this contribution can be interpreted through three mechanistic models. One possibility is that receptor binding directly activates the sperm for fusion, defined here as inducing a transition of the SPARK complex into a transient fusogenic state. This model predicts that activated sperm should be capable of fusing with eggs from different species or even with heterologous cells. However, it does not explain why soluble Bouncer inhibits fusion with wild-type eggs (*12*), disfavoring a sole role in activation.

A second scenario is that receptor binding instead primes SPARK for fusion. In this model, priming reflects receptor-induced changes in the SPARK complex, such as alterations in interfaces, affinities, or conformations, that render it more competent to transition into a fusogenic state. Consistent with this idea, AlphaFold modeling of the SPARK complex in the presence of Bouncer predicts conformational changes within SPARK (Fig. 5B), and co-immunoprecipitation experiments show increased association between Izumo1 and Spaca6 upon Bouncer binding (fig. S6A). Together, these observations suggest that Bouncer can induce changes in the SPARK complex prior to engagement with the egg membrane.

A third non-mutually exclusive possibility is that receptor binding bridges sperm for engagement with an additional egg membrane factor, which then primes or activates the fusion machinery. This model is consistent with the observation that soluble Bouncer inhibits fertilization in wild-type conditions (*12*) yet permits fusion in Bouncer-deficient eggs (Fig. 5C). This model assumes that Bouncer sequesters the additional membrane factor in wild-type eggs, which would remain accessible in the absence of Bouncer.

Taken together, we propose that under physiological conditions, Bouncer primes and/or bridges the SPARK complex to enable membrane fusion. This priming ability is reminiscent of viral systems, in which soluble receptors can modulate infection but require membrane-associated proteins for uptake and fusion (*70–72*). A similar functional framework, namely an opposing role of Bouncer in WT vs *bouncer-/-*eggs, may extend to mammals. Sperm pre-incubated with egg-derived CD9-containing vesicles can fuse with CD9-deficient eggs (*61*). In contrast, shedding of egg receptors such as JUNO and CD9 from WT eggs after fertilization has been proposed to limit polyspermy by acting as a sink for sperm in mice (*19*, *73*, *74*). Together, these observations raise the possibility that, despite molecular divergence, egg receptors across species couple sperm-egg binding to activation of the SPARK complex.

From an evolutionary perspective, our findings suggest that SPARK-mediated fertilization may have emerged from more ancient gamete fusion systems. The fusogen HAP2/GCS1, a member of the fusexin family, is widely conserved across plants, algae, protozoans, and some invertebrates, and fusexins have even been identified in archaea (*54–57*, *75*). Interestingly, DCST1/2 are among the most conserved components of the SPARK complex and are present in some invertebrates, partially overlapping with HAP2-containing lineages (Fig. 4, fig. S5G). This raises the possibility that ancestral fertilization mechanisms relied on HAP2-mediated fusion, which was later supplemented and/or replaced by DCST-based mechanisms (*46*, *76*). We speculate that this ancestral machinery subsequently expanded into a multi-protein assembly, incorporating additional factors such as TMDD1 and FAM187A, and later vertebrate-specific components. In this framework, the SPARK complex represents an evolutionary innovation that integrates ancient membrane-interacting proteins with newly evolved modules to enable tightly regulated, receptor-dependent sperm-egg fusion.

## Acknowledgements

We thank Vienna Biocenter Core Facilities (VBCF) for their expert support, in particular all members of the proteomics facility at IMP/IMBA/GMI using the VBCF instrument pool, members of the ProteinTechnology Facility for purifying recombinant soluble BIR2-Bouncer and BIR2, and all members of the IMP/IMBA/ GMI BioOptics Facility; the aquatics facility personnel from the IMP for taking excellent care of zebrafish, Olivia Füssl for help with genotyping transgenic fish, Sarah Liedl and Auwal Tanko for experimental help with zebrafish; Saki Nishioka, Kaito Yamamoto, and Chihiro Emori from the Ikawa lab for the generation of gene-manipulated mice; Keiko Murata from the Central Instrumentation Laboratory of Research Institute for Microbial Disease for performing Sanger sequencing on mouse PCR sample; the IMBA Fly and Worm Facility, in particular Peter Duchek and Joseph Gokcezade, for creating the *spe-40/fam187* mutant *C. elegans*; Jörg Fallmann for bioinformatic support; Amber Krauchunas and Steven Tang for sharing their pre-prints prior to submission to bioRxiv; Clemens Plaschka, Yongchan Lee, Tomohiro Nishizawa, Harmit Malik, Melody Campbell, Jessica Stock, Sascha Martens, Justyna Sawa-Makarska as well as the Vienna BioCenter Membrane Club for fruitful discussions; Angela Anderson (Life Science Editors), Diana Pinheiro, Clemens Plaschka and Theresa Zeisner for valuable feedback on the manuscript; and the entire Ikawa and Pauli labs for continuous discussions and support.

## Author contributions

V.E.D., J.P.S., and A.P. conceived the study; V.E.D. designed, performed, and analyzed fish experiments with contributions from K.P., A.Blaha, A.A.R.A., and M.M; J.P.S. performed and analyzed structural predictions of the SPARK complex; H.W., S.T. and Y.L. performed and analyzed mouse experiments with contributions from H.-Y.C. and T.N.; V.E.D., J.P.S., K.P. and A.Blaha helped in designing murine MS experiments and collecting samples for MS from murine sperm and testes; A.S. conducted phylogenetic analysis; J.A.H. conducted positive selection analysis; G.D. and M.N. performed the MS-based reanalysis of the Tmem81-IP and identified zebrafish Fam187a; K.S. and G.K. performed the PRM mass spectrometry experiments supervised by E.R.; H.M. performed the experiments in *C. elegans* supervised by A.Burga; A.P. supervised and coordinated the study, and M.I. supervised and coordinated the mouse research; V.E.D., J.P.S. and A.P. wrote the manuscript with contributions from all other authors.

## Materials and Methods

### Zebrafish and mouse research

Zebrafish (*Danio rerio*) were raised according to standard protocols with a 14/10h light/dark cycle at 28 °C water temperature. Wild-type zebrafish were acquired by crossing TL (Tupfel Longfin) and AB zebrafish. All zebrafish experiments were conducted with sexually mature males and females (at least 3 months-old) up to a maximum age of 1.5 years according to Austrian and European guidelines for animal research and approved by the Amt der Wiener Landesregierung, Magistratsabteilung 58—Wasserrecht (zebrafish protocols MA 58-221180-2021-16 and MA 58 – 357704-2026-12).

B6D2F1/Slc and Slc:ICR mice (Japan SLC, Inc.) were maintained under a 12-h light/dark cycle with ad libitum access to food and water. All experiments utilized male mice aged 8 weeks or older and female mice aged 7 weeks or older. Animals were euthanized via cervical dislocation following anesthesia. All experimental procedures were approved by the Animal Care and Use Committee of the Research Institute for Microbial Diseases, the University of Osaka (protocol #Biken-AP-R03-01), and were performed in accordance with institutional and national guidelines.

### AlphaFold predictions

Amino acid sequences of the mature proteins (excluding signal peptides and propeptides) for AlphaFold predictions were obtained from Uniprot. Signal peptide sequences were removed by predicting signal peptides with SignalP6.0 (*77*). The full-length sequences of zebrafish Tmdd1 and Fam187a were obtained as described below. To assess protein-protein interactions of known factors and novel candidates by PEAK scores (fig. S1A-B), an AlphaFold Multimer screen was conducted as previously described (*34*). In summary, a ‘baits’ file was generated with either the sequences of the zebrafish TIS trimer or the Dcst1/2 dimer. A ‘candidates’ file was created with the proteins that were significantly enriched in the co-IP/MS experiment from zebrafish sperm expressing Tmem81-3xFLAG (Fig. 1C). All combinations between bait and candidate proteins were predicted by AlphaFold Multimer. Structure predictions of individual proteins and protein complexes were conducted using the AlphaFold 3 server (*78*). All depicted models and their corresponding PAE plots are the top-scoring model generated by AlphaFold 3. Structure models and interaction networks (Fig. 1H) were visualized using ChimeraX-1.10 (*79*) and PAE plots were visualized using the PAE viewer webserver (*80*).

### Generation of zebrafish *tmdd1* and *fam187a* mutants

Zebrafish knockout fish for *tmdd1* and *fam187a* were generated by CRISPR/Cas9-mediated mutagenesis according to standard protocols. In brief, 1 guide RNA (gRNA) targeting the coding exons of *tmdd1* or *fam187a* was generated according to published protocols (*81*) by oligo annealing followed by T7 polymerase-driven *in vitro* transcription (gene-specific targeting oligos: tmdd1_gRNA: 5’-GAGCACTTCAACAGATCGGCAGG-3, fam187a_gRNA: 5’-CTACACGGGTCTGCTCACCCTGG-3’,common tracer oligo AAAAGCACCGACTCGGTGCCACTTTTTCAAGTTGATAACGGACTAGCCTTATTTTAACTTGCTATTTCTAGCTCTAA AAC). Cas9 protein and gRNAs were co-injected into the cell of one-cell stage TLAB embryos. Putative founder fish were outcrossed to TLAB wild-type fish. Founder fish carrying germline mutations were identified by size differences in the gene-specific amplicons in a pool of embryo progeny (primers: tmdd1_gt_F: 5’-AGCACTAAAGGACTGGCTTCAG-3’ and tmdd1_gt_R: 5’-TATTGATTGATGTGGTGCTGTG-3’; fam187a_gt_F: 5’-CCTCTGGTGAATTCTTCTACGG-3’ and fam187a_gt_R: 5’-GGAGCCTCTGTCTCTGAATCAC-3’). Embryos from founder fish were raised to adulthood. Heterozygous males and females of each mutant genotype were incrossed to generate homozygous mutant embryos. Sanger sequencing of homozygous mutant embryos identified the nature of the mutation. The *tmdd1* mutation is a 17-nt insertion in exon 3, which results in a frameshift mutation and a premature stop codon after 114 amino acids. The *fam187a* mutation is a 4-nt deletion, which results in a frameshift mutation and a premature stop codon after 189 amino acids. Genotyping of mutant fish was performed by PCR, and detection of the mutant product was performed by standard gel electrophoresis using a 4% agarose gel.

### Generation of zebrafish transgenic lines

Zebrafish transgenic lines expressing Tmdd1-mScarlet3 and Fam187a-3xFLAG-sfGFP under the control of the testis-specific *dcst2* promoter were generated by Tol2-mediated transgenesis (*tmdd1* and *fam187a* ORFs were PCR-amplified from cDNA derived from adult zebrafish testis and subcloned into a pMTB vector for Tol2 transgenesis). In brief, *tmdd1+/-* or *fam187a+/-* zebrafish embryos were injected at the 1-cell stage with 35 ng/mL *tol2* mRNA, 15 ng/mL of the respective expression construct and 0.083% phenol red solution. The embryos were kept in E3 medium at 28 °C, and those with strong somatic expression of the transgenesis markers were considered putative founders and raised to adulthood. Putative founder fish were crossed with *tmdd1+/-* or *fam187a+/-* zebrafish and the progeny was screened for the transgenesis marker indicating stable germline transmission. The transgenic progeny was raised to adulthood and genotyped to identify transgenic fish in a homozygous mutant (*tmdd1-/-* or *fam187a-/-*) background.

### Fertility assessment of adult zebrafish

Zebrafish *in vivo* fertilization rate measurements were performed by counting fertilization rates of natural matings. The evening prior to mating, the fish assessed for fertility and a TLAB wild-type fish of the opposite sex were separated in breeding cages. The next morning, the fish were allowed to mate. Eggs were collected and kept at 28 °C in E3 medium (5 mM NaCl, 0.17 mM KCl, 0.33 mM CaCl_2_, 0.33 mM MgSO_4_, 10^-5^% Methylene Blue). The rate of fertilization was assessed approximately 3 hours post-laying. By this time, fertilized embryos have developed to 1000-cell stage embryos, while unfertilized eggs resemble one-cell stage embryos.

### Sperm approach and sperm motility assays

To collect unactivated sperm, 2-4 wild-type, *tmdd1 -/-* and *fam187a -/-* male zebrafish were anesthetized with 50 mg/L tricaine for 10 min and subsequently anesthetized with 160 mg/L tricaine for 1-2 min. To assess sperm approaching the micropyle of wild-type zebrafish eggs; sperm was stored on ice in 100 µL of Hank’s saline containing 0.5 mM MitoTracker Deep Red FM (Molecular Probes). Unactivated, mature eggs were expelled into a dry petri dish by gentle pressure on the belly of an anesthetized wild-type female. To prevent activation, eggs were kept in sorting medium (Leibovitz’s medium, 0.5% BSA, pH 9.0) at RT. Eggs were transferred to a Petri dish with cone-shaped agarose molds (1.5% agarose in E3 medium) filled with sorting medium and oriented to allow visualization of the micropyle and the dish mounted on a microscope. Sorting media was removed and carefully replaced with 1 mL of E3 medium. Approximately 250,000 – 400,000 stained sperm were added as close as possible to the eggs. Time-lapse images were acquired using a Zeiss LSM800 Examiner Z1 upright microscope equipped with a 20x/1.0 Plan-Apochromat water-dipping objective. Image sequences were analyzed using Fiji (*82*).

Sperm motility was assessed as previously described (*12*). Sperm was isolated from anesthetized wild-type, *tmdd1 -/-* and *fam187a -/-* male fish, collected in 100 µL of Hank’s saline, and kept on ice. Sperm were activated by adding E3 medium in a 1:4 dilution (sperm:E3 medium). Immediately after activation, approximately 10.000 – 20.000 sperm cells were loaded into a 10 µm thick chamber slide (Leja counting chamber, SC 10–01–04-B). Sperm motility was recorded for 1 min, starting no later than 30 sec post-activation, using a Zeiss Axio Imager.Z2 microscope equipped with a 10x/0.3 Plan-Neofluar objective using darkfield illumination. Sperm tracks were analyzed using Fiji with the ‘‘TrackMate’’ plugin (*82*, *83*). Sperm that were present in the movie for the first 100 timeframes after sperm activation were tracked. Sperm heads were marked using the LoG Detector algorithm and linked across frames through the Advanced Kalman Tracker to create a track. Each track was manually verified, and tracking errors were corrected using the TrackScheme tool. Track data was exported to obtain the average sperm velocity.

### Sperm-egg binding assay

Sperm-egg binding assays using activated, dechorionated eggs were performed as previously described (*12*). To collect unactivated sperm, wild-type, *tmdd1 -/-* and *fam187a -/-* male zebrafish were anesthetized with 50 mg/L tricaine for 10 min and subsequently anesthetized with 160 mg/L tricaine for 1-2 min. Unactivated sperm was collected from 2-6 fish per genotype and kept in 100 µL of Hank’s saline containing 0.5 µM MitoTracker Deep Red FM (Molecular Probes) on ice. Unactivated, mature eggs were collected from a wild-type female fish, activated by the addition of E3 medium and incubated for 1 h. Subsequently, 1–2 eggs were manually dechorionated using forceps and transferred to an imaging dish with cone-shaped agarose molds, filled with E3 medium and then mounted on a microscope. After focusing on the egg plasma membrane, the objective was briefly raised and approximately 250,000 – 400,000 labeled sperm cells were added. Imaging was performed with a Zeiss LSM800 Examiner Z1 upright system using a 20x/1.0 Plan-Apochromat water-dipping objective. Images were acquired until sperm were no longer motile (approximately 5 min). For analysis of sperm-egg binding, the number of stably bound sperm was quantified. Sperm were counted as bound when they remained in the same position for at least 1 min following a 90 sec activation and approach time window. Data was plotted as the number of sperm bound per 100 µm of egg membrane for one minute.

### Zebrafish sperm and egg collection for *in vitro* fertilization assays

Zebrafish unactivated gametes were collected as previously described (*12*). In brief, the evening prior to sperm and egg collection, male and female zebrafish were separated in breeding cages (one male and one female per cage). To collect unactivated sperm, wild-type and *tmdd1 -/-* male zebrafish were anesthetized with 50 mg/L tricaine for 10 min and subsequently anesthetized with 160 mg/L tricaine for 1-2 min. Unactivated sperm was collected in a capillary by mouth pipetting under a dissecting microscope. The sperm was stored on ice in Hank’s saline solution (5.4 mM KCl, 0.137 M NaCl, 1 mM MgSO_4_, 4.2 mM NaHCO_3_, 0.25 mM Na_2_HPO_4_, 1.3 mM CaCl_2_). For BIR2 or BIR2-Bouncer IVFs, 5 µM recombinant protein was added to sperm in Hanks (final sperm concentration: 75,000 sperm/µL) and incubated for >20 min on ice. To collect unactivated eggs, *bouncer -/-* female zebrafish were anesthetized as described, and eggs were expelled into two dry petri dishes by gentle pressure on the belly of the female. 45 µL of sperm solution (3-4 million total sperm) pre-incubated with BIR2 or BIR2-Bouncer was added to eggs that were subsequently activated by the addition of 500 µL of E3 medium (5 mM NaCl, 0.17 mM KCl, 0.33 mM CaCl_2_, 0.33 mM MgSO_4_, 10^-5^% Methylene Blue). After 3-5 min, Petri dishes were filled with E3 medium and kept at 28 °C. The rate of fertilization was assessed approximately 3 hours post-laying.

### Preparation of zebrafish sperm samples for co-IPs and PRM

The Tmem81-3xFLAG-sfGFP zebrafish sperm co-IP (Fig. 1C) was performed in a previous study (*12*). For the Spaca6-3xV5 zebrafish sperm co-IP (fig. S1E), 4 - 7 x 10^8^ sperm were collected in Hank’s saline. Samples were crosslinked with formaldehyde (final concentration of 1% formaldehyde (w/v); note that the bottle of formaldehyde used for this IP experiment was opened 2-month prior to the experiment, which decreased its cross-linking activity) and incubated for 10 min at room temperature. 100 mM Tris-HCl (pH 8) was added and incubated for 10 min to quench the crosslinking reaction. Prior to lysis, sperm was pelleted by spinning samples at 2000 g for 3 min at 4 °C and resuspended in about 30 µL Hank’s saline. The pellet was then resuspended in 200 µL lysis buffer (15 mM NaHEPES pH 7.3, 30 mM NaCl, 1 mM MgCl_2_, 1% Brij35, 1 U/mL benzonase (Merck), 1X complete protease inhibitor (EDTAfree, Roche), 0.5% (w/v) SDS) and incubated on ice for 2 hours. Insoluble debris was pelleted at 21,000 g at 4 °C for 10 min. The supernatant was used for the immunoprecipitation with V5-Trap Magnetic Beads (Chromotek), which was performed overnight at 4 °C and shaking at 1100 rpm. The beads were washed two times with lysis buffer and three times with wash buffer (15 mM HEPES, 30 mM NaCl) before being transferred to a clean tube to remove any traces of detergent.

For the co-IP from Spaca6-3xV5 zebrafish sperm incubated with recombinant BIR2 or BIR2-Bouncer (fig. S6A), 0.75 – 1.4 x 10^9^ sperm were collected in Hank’s saline. Samples were split into two and were incubated with 88.5 µg BIR2-1xSTREP or 125 µg BIR2-Bouncer-2XSTREP for 45 min on ice. Prior to lysis, sperm was pelleted by spinning samples at 800 g for 3 minutes at 4 °C and resuspended in about 30 µL Hank’s saline. The pellet was then resuspended in 150 µL lysis buffer (15 mM NaHEPES pH 7.3, 30 mM NaCl, 1 mM MgCl_2_, 1% Brij35, 1 U/mL benzonase (Merck), 1X complete protease inhibitor (EDTAfree, Roche), 0.2% (w/v) SDS) and incubated on ice for 2 hours. Insoluble debris was pelleted at 21,000 g at 4 °C for 10 min. The supernatant was used for the immunoprecipitation with V5-Trap Magnetic Beads (Chromotek) overnight at 4 °C and shaking at 1100 rpm. The beads were washed two times with lysis buffer and three times with wash buffer (15 mM HEPES, 30 mM NaCl) before being transferred to a clean tube to remove any traces of detergent.

Beads were resuspended in 40 µL of 100 mM ammonium bicarbonate (ABC) supplemented with 400 ng of lysyl endopeptidase (Lys-C, Fujifilm Wako Pure Chemical Corporation) and incubated for 4 h on a thermo-shaker with 1200 rpm at 37 °C. The supernatant was transferred to a fresh tube and reduced with 0.5 mM Tris 2-carboxyethyl phosphine hydrochloride (TCEP, Sigma) for 30 minutes at 60 °C and alkylated in 4 mM methyl methanethiosulfonate (MMTS, Fluka) for 30 min at room temperature. Subsequently, the sample was digested with 400 ng trypsin (Trypsin Gold, Promega) at 37 °C overnight. The digest was acidified by addition of trifluoroacetic acid (TFA, Pierce) to 1%.

### Generation of a mouse TMEM81-3xFLAG transgenic line

To produce a transgenic TMEM81-3xFLAG-expressing mouse line, the ORF of *Tmem81* was cloned from mouse testis cDNA by PCR and inserted into a plasmid encoding an *Izumo1* promoter and a rabbit beta-globin polyadenylation (polyA) signal. The transgene, along with the promoter and polyA signal, was digested by restriction enzymes and was microinjected into the pronuclei of zygotes obtained from mating between homozygous *Tmem81* KO females and heterozygous *Tmem81* KO males (*12*). The injected zygotes were cultured in the KSOM medium until the two-cell stage and transplanted into the oviductal ampulla of 0.5 day pseudo-pregnant ICR females. Founder animals were obtained by natural delivery or Cesarean section after 19 days of pregnancy. The transgene was identified by PCR using primers targeting the *Izumo1* promoter and the polyA signal. Frozen spermatozoa from *Tmem81*^+/–^ transgenic males (B6D2-Tg(*Izumo1-Tmem81/ 3xFLAG*)1Osb) have been deposited to the Riken BioResource Center (RIKEN BRC; web.brc.riken.jp/en) and the Center for Animal Resources and Development, Kumamoto University (CARD R-BASE; <u>cardb.cc.kumamoto-u.ac.jp/transgenic</u>).

### Generation of mouse *Tmdd1* and *Fam187a* mutants

*Tmdd1* and *Fam187a* knockout (KO) mouse lines were generated using CRISPR/Cas9-mediated genome editing. To delete the coding region, two single-guide RNAs (sgRNAs) were designed to flank either the *Tmdd1* or the *Fam187a* locus. B6D2F1/Slc female mice were superovulated via intraperitoneal injection of CARD HyperOva (Kyudo Co.) followed by human chorionic gonadotropin (hCG; ASKA Pharmaceutical Co. Ltd.) 48 hours later. These females were mated with wild-type B6D2F1/Slc males, and two-pronuclear (2PN) zygotes were subsequently collected from the oviducts. Ribonucleoprotein (RNP) complexes, consisting of CRISPR RNA (crRNA), trans-activating crRNA (tracrRNA), and Cas9 protein, were introduced into the 2PN zygotes using a NEPA21 super electroporator (NEPA GENE). Electroporated zygotes were cultured in potassium simplex optimization medium (KSOM) until the two-cell stage. Surviving embryos were then transferred into the oviductal ampulla of 0.5-day pseudopregnant Slc:ICR females. Founder mice were obtained through natural delivery or Cesarean section. Initial screening of founder mice was performed by PCR genotyping, followed by Sanger sequencing to characterize the specific mutant alleles. Established lines were maintained by backcrossing with wild-type B6D2F1/Slc mice. Frozen sperm from *Tmdd1^+/-^* and *Fam187a^+/-^* males will be available through the RIKEN BioResource Center (RIKEN BRC) and the Center for Animal Resources and Development (CARD), Kumamoto University.

### Mouse mating tests

To assess *in vivo* fertility, sexually mature *Tmdd1^-/-^*and *Fam187a^-/-^* male mice were housed individually with three sexually mature wild-type females for a period of four to eight weeks. Successful copulation was confirmed by the daily monitoring of vaginal plugs. The number of pups was recorded at birth. Following the mating period, male mice were removed, and females were monitored for an additional three weeks to ensure the delivery and documentation of any final litters resulting from the last mating events.

### Mouse sperm motility

Cauda epididymal sperm were extracted from adult male mice under a dissection microscope using micro spring scissors and fine forceps. The extracted sperm were immediately dispersed into 100 µL drops of Toyoda-Yokoyama-Hoshi (TYH) medium. Sperm motility was quantified using the CEROS II sperm analysis system (Hamilton Thorne Biosciences) after 10 min and 120 min of incubation in TYH medium at 37 °C under 5% CO_2_.

### Mouse ZP-free IVF

Cauda epididymal sperm from sexually mature *Tmdd1^-/-^* and *Fam187a^-/-^* males were preincubated in TYH medium for 2 h at 37 °C under 5% CO_2_ to induce capacitation. Cumulus-oocyte complexes (COCs) were recovered from the oviductal ampulla of superovulated wild-type females and transferred into fresh TYH drops. To generate zona-free oocytes, 1 mg/mL collagenase (Sigma-Aldrich) was added to the drops to facilitate the removal of both cumulus cells and the zona pellucida (ZP). These ZP-free oocytes were then inseminated with capacitated sperm at a final concentration of 2 × 10^4^ sperm/mL. Fertilization success was evaluated 6 hours post-insemination by the presence of two pronuclei (2PN) using a phase-contrast microscope.

### Mouse sperm-oocyte binding and fusion assay

To evaluate oocyte binding and fusion ability, cauda epididymal sperm were preincubated in TYH medium for 1 h. To monitor the acrosome reaction, sperm were then incubated with a rat anti-IZUMO1 monoclonal antibody (*84*), together with an Alexa Fluor 488-conjugated anti-rat IgG secondary antibody (Invitrogen, A11006) in TYH medium for an additional 1 h. Simultaneously, ZP-free oocytes were prepared from superovulated wild-type females via collagenase treatment. To facilitate the identification of fused sperm nuclei, ZP-free oocytes were stained with Hoechst 33342 (1:10000) for 15 min and washed thoroughly in fresh TYH medium. The ZP-free oocytes were inseminated with the sperm at a density of 2 × 10^5^ sperm/mL. After a 30-min incubation, sperm-oocyte complexes were fixed in 0.1% paraformaldehyde (PFA) in flushing holding medium (FHM) for 15 min and washed gently. The number of acrosome-reacted sperm (IZUMO1-positive) bound to the oocytes, and the number of fused sperm (Hoechst 33342-positive) were quantified using a BZ-X810 fluorescence microscope (Keyence).

### Preparation of mouse sperm samples for co-IP

For co-IP from mouse sperm expressing TMEM81-3xFLAG, epididymal sperm from 5-7 mice per replicate and condition (WT, negative control or TMEM81-3xFLAG) was isolated. Sperm samples were lysed for 2 h at 4 °C in 100 μL IP lysis buffer (50 mM Tris pH 7.5, 150 mM NaCl, 10% glycerol, 1% TritonX-100, protease inhibitor cocktail (nacalai tesque, 25955-11)). Insoluble material was pelleted by centrifugation at 15,000 rcf for 15 min at 4 °C. For the IP, FLAG M2 antibody (Sigma, F3165) was crosslinked to dynabeads using the Pierce Crosslink IP kit (Thermo Scientific, 26147) at a concentration of 0.4 μg antibody per μL bead slurry. Sperm lysates were added to anti-FLAG crosslinked bead slurry and incubated o/n at 4 °C. The amount of beads used was normalized to 3 Mio sperm cells per μL of bead slurry. Beads were washed three times in 500 μL IP lysis buffer, transferred to fresh tubes and washed six times in 500 μL detergent-free IP lysis buffer (50 mM Tris pH 7.5, 150 mM NaCl).

Beads were resuspended in 40 µl of 100 mM ammonium bicarbonate (ABC), supplemented with 400 ng of lysyl endopeptidase (Lys-C, Fujifilm Wako Pure Chemical Corporation) and incubated for 4 hours on a Thermo-shaker with 1200 rpm at 37 °C. The supernatant was transferred to a fresh tube and reduced with 0.5 mM Tris 2-carboxyethyl phosphine hydrochloride (TCEP, Sigma) for 30 minutes at 60 °C and alkylated in 4 mM methyl methanethiosulfonate (MMTS, Fluka) for 30 min at room temperature. Subsequently, the sample was digested with 400 ng trypsin (Trypsin Gold, Promega) at 37 °C over night. The digest was acidified by addition of trifluoroacetic acid (TFA, Pierce) to 1%.

### Sample preparation of sperm and testis lysates used for DIA-based shotgun and targeted LC-MS/MS

For the measurement of relative protein abundance, 2.5 - 3.5 x 10^7^ zebrafish sperm per biological replicate were collected and lysed in SDT buffer (4% SDS, 100 mM DTT, 100 mM Tris-HCl pH 7.5, 1 mM MgCl_2_, 7 U/mL benzonase (Merck)). Mouse sperm was isolated from the epididymis of one mouse per biological replicate and lysed in SDT lysis buffer. Mouse testis samples (from one mouse per genotype per replicate) were dissected in PBS and subsequently lysed and homogenized in SDT buffer. For lysis, samples were incubated for 10 min at 95 °C before sonication with an ultrasonication probe connected to an Ultrasonic processor UP100H (Hielscher) with 30 cycles (0.5 s, 50% amplitude). The protein concentration was determined using the Pierce 660-nm Protein Assay Reagent kit (PN 22660; Thermo Fisher Scientific) with Ionic Detergent Compatibility Reagent (PN 22663; Thermo Fisher Scientific).

The protein extract was mixed with 200 µl of 8 M urea in 0.1 M TEAB pH 8.5 and processed in Microcon filter units with 30 kD cut-off using the FASP method as described (*85*), using trypsin (Mass spectrometry grade; Promega) in a ratio of 1:20 (protease:protein; w/w). The final peptide concentration was determined by separating an aliquot of each sample on a LC-UV system equipped with monolithic trap (PepSwift monolithic PS-DVB trap (100 mm x 5 cm), Thermo Fisher Scientific) and analytical column (200 mm x 5 cm, Thermo Fisher Scientific) and measuring the absorption at UV 214 nm. The integrated peak area was quantified by relating to the peak area of 100 ng of Pierce HeLa protein digest standard (PN 88329; Thermo Fisher Scientific). The peptide solution was frozen at -80 °C before further processing.

### Assembly of a custom ORF database for zebrafish testis

To build a more complete testis-specific MS reference that captures transcript diversity beyond standard annotations, we leveraged the available deeply sequenced paired-end testis RNA-seq data available under GEO number GSE111882. We used these data to perform reference-guided transcriptome assembly, by reprocessing the pooled testis sample reads with the nf-core/rnaseq workflow (v3.17.0) enabling read assembly with option --stringtie_ignore_gtf and providing genome and gene annotation from *Danio rerio* Ensembl release 113 as reference (TrimGalore v0.6.10; STAR v2.6.1d; StringTie v2.2.3; GRCz11 assembly). Coding regions within assembled transcripts were predicted by TransDecoder v5.7.1 setting the minimum protein length to 40 (TransDecoder.LongOrfs -m 40).

### Re-analysis of Tmem81 zebrafish sperm co-IP/MS proteomics data

For the identification of unannotated protein-coding genes, proteomics measurements were searched as previously described (*12*) against the Ensembl reference proteome as well as a custom database of ORFs (208.551 sequences, 29.296.085 residues) that was generated as described above. Raw MS data were loaded into Proteome Discoverer (PD, version 2.5.0.400, Thermo Fisher Scientific). All MS/MS spectra were searched using MSAmanda v2.5.0.19924. Peptide-spectrum matches from those two searches were intersected at the peptide level, and only RNA-seq–derived ORFs supported by at least two PSMs corresponding to peptides not identified in the reference proteome were retained. A final combined database search was then performed using both the Ensembl proteome and the filtered RNA-seq ORFs (195 sequences, 135.545 residues) as well as the contaminant database and four proteins of interest (Tmem81, Spaca6, Flag-GFP, Fam187a, 1.277 residues), enabling the identification of known proteins alongside previously unannotated protein-coding candidates.

### DIA LC-MS/MS measurements

The Vanquish Neo UHPLC-System was coupled to an Orbitrap Astral mass spectrometer equipped with a Nanospray Flex ion source (all parts Thermo Fisher Scientific).

Peptides were loaded onto a trap column (PepMap C18, 5 mm × 300 μm ID, 5 μm particles, 100 Å pore size, Thermo Fisher Scientific) using 0.1% TFA as mobile phase. After loading, the trap column was switched in line with the analytical column (Aurora Ultimate C18 25cm × 75 μm ID, 1.7 μm particles, 120 Å, operated at 50 °C, Ionopticks). Peptides were eluted using a flow rate of 300 nl/min, starting with the mobile phases 98% A (0.1% formic acid in water) and 2% B (80% acetonitrile, 0.1% formic acid) and linearly increasing to 35% B over the next 60 min followed by an increase to 95% B in 1.7 min, a 4 min hold at 95% B, and re-equilibration with 2% B for three column volumes.

The Orbitrap Astral mass spectrometer was operated in data-independent mode, performing a full scan in the Orbitrap every 0.6 sec (*m/z* range 380–980; resolution 240,000; AGC target 1,000,000, maximum injection time 5 msec). MS/MS spectra were acquired in the Astral analyzer by isolating 4 Da windows (5 Da for full lysates) across 380–880 *m/z* or 380-980 *m/z* (for full lysates) and fragmenting precursor ions with HCD collision energy of 25% with a maximum injection time of 5 msec (3 msec for full lysates) or until an AGC target of 30,000 was reached. Fragment ions ranging from 150–2000 *m/z* were acquired.

### DIA Proteomics data analysis

DIA data was analysed in Spectronaut (*86*) (Biognosys). Trypsin/P was specified as a proteolytic enzyme and up to 2 missed cleavages were allowed in the Pulsar directDIA+ search. Dynamic mass tolerance was applied for calibration and main search. The search was performed against the Ensembl zebrafish database (v111, 30.135 entries) or the Uniprot mouse database (20260203, 21.852 entries), with common contaminants and tags appended. Carbamidomethylation or beta-methylthiolation of cysteine was searched as fixed modification, whereas oxidation of methionine and acetylation at protein N-termini were defined as variable modifications. Peptides with a length between 7 and 52 amino acids were considered and results were filtered using Spectronaut default filtering criteria (Precursor Qvalue<0.01, Precursor PEP<0.2, Protein Qvalue <0.01 per Experiment and <0.05 per Run, Protein PEP<0.75). Quantification was performed as specified in Biognosys BGS Factory Default settings, grouping Peptides by Stripped Sequence and performing protein inference using IDPicker. Cross-Run Normalization in Spectronaut was deactivated due to subsequent mode normalization.

Spectronaut results were exported using Pivot Reports on the Protein and Peptide level and converted to Microsoft Excel files using our in-house software MS2Go [https://ms.imp.ac.at/?action=ms2go]. For DIA data MS2Go utilizes the python library msReport (developed at the Max Perutz Labs Proteomics Facility) for data processing. Abundances were normalized by the mode of protein ratios in msReport and missing values were imputed with values obtained from a log-normal distribution with a mean of 100. To compensate for different protein lengths, protein quantification was then normalized using iBAQ (*87*). Statistical significance of differentially expressed proteins was determined using limma (*88*).

### Targeted proteomics LC-MS/MS analysis

Nano-LC-MS/MS analysis was performed on a nano-HPLC system (Vanquish Neo UHPLC-System) operated as described above, applying a 90 min gradient from 2 to 35% B. The Orbitrap Exploris 480 mass spectrometer was operated by a mixed MS method, which consisted of one full scan (*m/z* range 380-1.500; 15.000 resolution; target value 1E6) followed by the PRM of targeted peptides from an inclusion list (isolation window 0.8 *m/z*; normalized collision energy (NCE) 32; 30.000 resolution, AGC target 2E5). The maximum injection time was set to 800 ms. For method development and validation of the peptides of interest, stable isotope-labeled synthetic peptides (13C-labeled Lysine and Arginine) of the proteins of interest were synthesized. Additionally, peptides of proteins considered to show constant expression levels (Uniprot IDs for ‘control proteins’ for zebrafish sperm: Q6P4P4, F6P3L9, Q6PEI6, B8JI08, Q7SXA3, A0A2R9YJL1; Uniprot IDs for ‘control proteins’ for mouse sperm and testis: P00342, P68134, P32037, O70325) were analyzed by PRM and used for normalization of total protein amounts. At least four unique peptides of the proteins of interest were measured per protein. At least three biological replicates were measured for all sperm or testis samples of each genotype. All peptides included in the final sPRM method are provided in Table S1. A scheduled PRM method (sPRM) development, data processing and manual evaluation of results were performed in Skyline (Skyline-daily 64-bit, 25.1.1.258) (*89*). Spectra of unique peptides of the proteins of interest (zebrafish: Izumo1, Spaca6, Tmem81, Fam187a, Dcst1, Dcst2, Tmdd1; mouse: DCST1, DCST2, FIMP, FAM187A, IZUMO1, TMEM95, TMEM81, SPACA6, TMDD1) and the ‘control proteins’ were recorded. The sum of the peptide areas of each normalization protein was divided by the average sum across samples. The values were averaged across all normalization proteins to obtain a normalization factor for each sample that served as correction for total protein input. The sum of peptide areas of each protein of interest was scaled by the normalization factor for each sample. The relative protein abundance in each sample was determined by dividing by the average value of the wild-type samples.

### Evolutionary analysis

Orthologous sequences of Tmdd1 were collected in a Blast search with human TMDD1 (NCBI protein: NP_001373666.1, UniProt: P0DPE3) as query, in the UniProt reference proteomes or NCBI nr protein databases, applying significant E-value thresholds below 0.001 (blast+ v.2.8.1; (*90*), UniProt,Sayers 2024]). Hits were found in all vertebrate lineages, selected sequences aligned with MAFFT (-linsi mode,v7.525) (*91*), and visualized with Jalview v. 2 (*92*). Protein domains were annotated with HMMER (v.3.4) and the PFAM database v.38.2 (*93*) as well as interproscan in InterPro v. 108, (*94*). TMDD1 proteins are characterized by a signal peptide (*Homo sapiens* TMDD1: 1-27), a highly conserved extracellular DEATH domain (amino acids 26-167), a transmembrane region (amino acids 215-244) and a weakly conserved cytoplasmic tail. Iterative Psi-blast searches expanded the TMDD1 family to invertebrates, such as lancets, mollusks, segmented worms, and tardigrades. The DEATH domain of human TMDD1 hit to the gastropod *Pomacea canaliculata* TMDD1 (UniProt: A0A2T7PE00) in round 2 with a highly significant E-value of 2.38e-09, whereas other DEATH domain-containing families, such as ANK2, were hit in higher iterations (>round 7). The reciprocal Psi-blast with *Pomacea canaliculata* TMDD1 hit to *Homo sapiens* TMDD1 in round 2 with an E-value of 1.64e-06 and incorporates other DEATH domain proteins in round 5. The orthologous relationship of the vertebrate and invertebrate TMDD1 families was further supported in a phylogenetic tree calculated with IQTREE v.3.0.1 (*95*), that included other DEATH domain proteins as outliers. For comparison of extracellular and intracellular DEATH domain proteins, the PFAM DEATH domain was used in an HMM search in selected vertebrate UniProt reference proteomes. DEATH domains with a significant E-value of 1E-5 (full sequence and domain) were extracted with a BioPerl script. Tmdd1 DEATH domains were not detected with this restrictive E-value threshold and added to the sequence set. 456 regions were initially aligned with mafft, later on filtered for completeness and sequence errors in Jalview, leading to a final set of 366 regions for the inference of a phylogenetic tree with IQTREE, with standard model selection using ModelFinder (*96*), ultrafast bootstrap (UFBoot2) support values (*97*) and the SH-aLRT test (*98*). The tree was visualized in iTOL v7 (*99*).

The FAM187 protein family was collected in a blast search starting with human FAM187A (UniProt: A6NFU0), applying the significant E-value threshold of 0.001. One hit was to the nematode *Plectus sambesii* Ig-like domain-containing protein (UniProt: A0A914WMH3) with an E-value of 1.71e-05, which was further on used to identify *Caenorhabditis elegans* Ig-like domain-containing protein CELE_Y37E11AR.7 with an E-value of 2e-20. FAM187 proteins are characterized by a signal peptide (*Homo sapiens* FAM187A: amino acids 1-17), an extracellular region with two Ig-like domains (amino acids 48-129, 273-361), a transmembrane region (amino acids 382-405) and a short cytoplasmic region. There are two paralogs in vertebrates, FAM187A and FAM187B, and a phylogenetic tree with IQTREE demonstrated that human FAM187A and FAM187B are co-orthologous to *Caenorhabditis elegans* CELE_Y37E11AR.7. This *C. elegans* protein was thus annotated as *C. elegans* Fam187, and is also referred to as Spe-40 (*45*). Only one Fam187 protein was identified in the transcriptome of *Danio rerio,* and it is closer to human FAM187A.

To determine the taxonomic distribution of fertilization factors, orthologous sequences of the respective protein families were selected with a broad taxonomic range, derived from sensitive HMM searches with family-specific domain models (from PFAM) or reciprocal blast searches. These reference sequences were used for blast searches in the UniProt reference proteomes or NCBI nr protein databases. An E-value threshold of 0.001 was applied to prevent collecting hits from protein families with similar domains. For the IZUMO1 protein family, the E-value threshold was set to 1e-10 to avoid inclusion of other Ig-like proteins. The taxonomic tree was generated with the NCBI Taxonomy Common Tree tool (*100*). The visualization was performed in iTOL v7 (*99*). For the UpSet plot, we only considered sequences from eukaryotic reference proteomes in UniProt, omitting vertebrates. For the visualization, we used python code generated by <u>Abacus.AI</u>. The reference sequence sets of each protein family are provided on request.

### Positive selection analysis

Positive selection analyses were performed as previously published (*101*). Coding sequences from all SPARK components as well as CD9, CD81, and JUNO in all *Murinae* species with annotated genomes in NCBI (*102*) were collected and codon-aligned via webPRANK (*103*), and manually trimmed of gaps that contribute noise to analyses. Alignments were used to generate phylogenetic trees via phyML (*98*). Gene-wide and site-specific positive selection was assessed via a streamlined wrapper (https://github.com/jayoung/pamlWrapper) for the codeml package in PAML (*104*). Whole gene dN/dS was calculated via model 0. For gene-wide analysis, the likelihoods of dN/dS fitting a model where either positive selection was (model 2) or was not (model 1) allowed were assessed; the same approach was subsequently used for site-specific positive selection by comparing model 8 to model 8a; we restricted classifying residues under positive selection to those with a Bayes Emperical Bayes posterior probability greater than 0.9 under model 8. Codon alignments can be found in Supplemental Data 1, and PAML analysis results in Table S2. Residues under positive selection were mapped onto the predicted SPARK complex structure and visualized using UCSF ChimeraX.

### Generation of a C. elegans spe-40/fam187 mutant

CRISPR/Cas9 genome editing in *C. elegans* was performed following previously described protocols (*105*). In brief, 250 ng/µl Cas9 protein (IDT) was incubated with 200 ng/µl of each crRNA (IDT) and 333 ng/µl tracrRNA (IDT) at 37 °C for 10 min to assemble ribonucleoprotein complexes. Subsequently, 2.5 ng/µl of the co-injection marker plasmid pCFJ90-mScarlet-I was added to the injection mix. For homology-directed repair, the donor oligo (IDT) was included at a final concentration of 200 ng/µl. Following injections into young hermaphrodites, mScarlet positive F1 progeny were singled, and their offspring screened by PCR and Sanger sequencing to detect successful editing. To generate a knockout allele of Y37E11AR.7 (*spe-40/fam187*), two crRNAs were designed to target exon 4 and exon 6 of the locus. Simultaneous cleavage at these sites resulted in a 634-bp deletion after repair, introducing a premature stop codon at position Ala84. Three independent alleles carrying the desired mutation were recovered. In all cases, hermaphrodites homozygous for the mutant allele were sterile. One allele was selected for further characterization and maintained in a heterozygous state: strain INK1307, Y37E11AR.7(abu575[p.Ala84*]/+) IV.

Sequences of the cRNAs used: crRNA1 5’-CTTCCAACTGATCCAGCCGA-3’ and crRNA2 5’-CGGATCGTGGA-GAGCAAACT-3’. Sequence of the Y37E11AR.7 (*spe-40/fam187*) repair oligo: 5’-TAGTGTCTCCCCGGAAGAC-GGTGTGGAAATTGATTCAACGTACAAGCCGTCG-TAACTGACTAACTAGGACCATAAAGTGCTATGTTGAGGTAAAGAAGTTGTAATGGTGTCAGCGG-3’. For genotyping, the following primers were used: Forward primer 5ʹ-CGAGACATCGGAGAGGAGAGCAG-3ʹ and Re-verse primer 5ʹ-GGAGGTAGTCGAGGGTTCTGATG-3ʹ. PCR amplification produced a 1320-bp fragment for the wild-type allele and a 670-bp fragment for the mutant allele.

### Characterization of the fertility defect of the *spe-40/fam187 C. elegans* mutant

Nematodes were grown on modified nematode growth medium (NGM) plates with 1% agar/0.7% agarose 20 °C. Plates were seeded with OP50 bacteria. Some of the strains used in this study were provided by the CGC, funded by the NIH Office of Research Infrastructure Programs (P40 OD010440). To characterize fertility defects associated with the *spe-40/fam187* mutation, brood size assays were performed. Starting from heterozygous mothers, 20 F1 L1 progeny were singled onto individual NGM plates. Once animals reached young adulthood, the number of laid embryos that developed normally was counted every 24 hours for four consecutive days. After the assay, the genotype of each mother was determined by PCR. No difference in total brood size was observed between homozygous wild-type (+/+) animals (n = 2) and heterozygous *spe-40/fam187* mutants (+/–) (n = 13). In contrast, 100% of homozygous mutant hermaphrodites (–/–) (n = 5) were completely sterile. Examination of plates containing homozygous mutants revealed large clusters of laid unfertilized oocytes. To determine whether sterility in *spe-40/fam187* (–/–) animals resulted from defective sperm, mating rescue experiments were performed. Six homozygous mutant adult hermaphrodites were crossed with wild-type N2 males. Eggs laid were counted every 24 hours for four days. In all cases, fertility was restored, with mutant hermaphrodites producing between 100-277 developing embryos.

### Data availability statement

The original contributions presented in the study are included in the article/supplementary material; mass spectrometry proteomics data will be deposited to the ProteomeXchange Consortium via the PRIDE partner repository; all inquiries should be directed to the corresponding authors.

## Supplementary Tables, Data and Movies

**Supplementary Table S1:** Analyzed mass spectrometry tables for shotgun and parallel reaction monitoring mass spectrometry experiments including re-analysis of zebrafish Tmem81 IP with peptide coverage and sequences of both Ensembl and testis RNA-seq based ORFs.

**Supplementary Table S2:** An expanded table of the *Murinae* positive selection analysis presented in fig. S5D. Log likelihood values were used for chi-square statistical comparisons between models allowing or not allowing for positive selection.

**Supplemental Data 1:** A codon alignment file for *Murinae* SPARK positive selection analysis.

**Supplementary Movie 1:** AlphaFold predictions of zebrafish, murine and human SPARK complex.

**Supplementary Movie 2:** Sperm-egg binding assay in zebrafish. While wild-type sperm can bind to wild-type eggs (left), *tmdd1* KO (middle) and *fam187* KO (right) sperm fail to bind.

**Supplementary Movie 3:** AlphaFold model (morph) of the predicted conformational change induced by Bouncer binding to the SPARK complex.

**Supplementary Movie 4:** DCST1/2 are predicted to adopt a bent conformation within the SPARK complex in which their hydrophobic tip is shielded through interaction with the N-terminus of TMEM81.

## Supplementary Figures

**Fig. S1.**
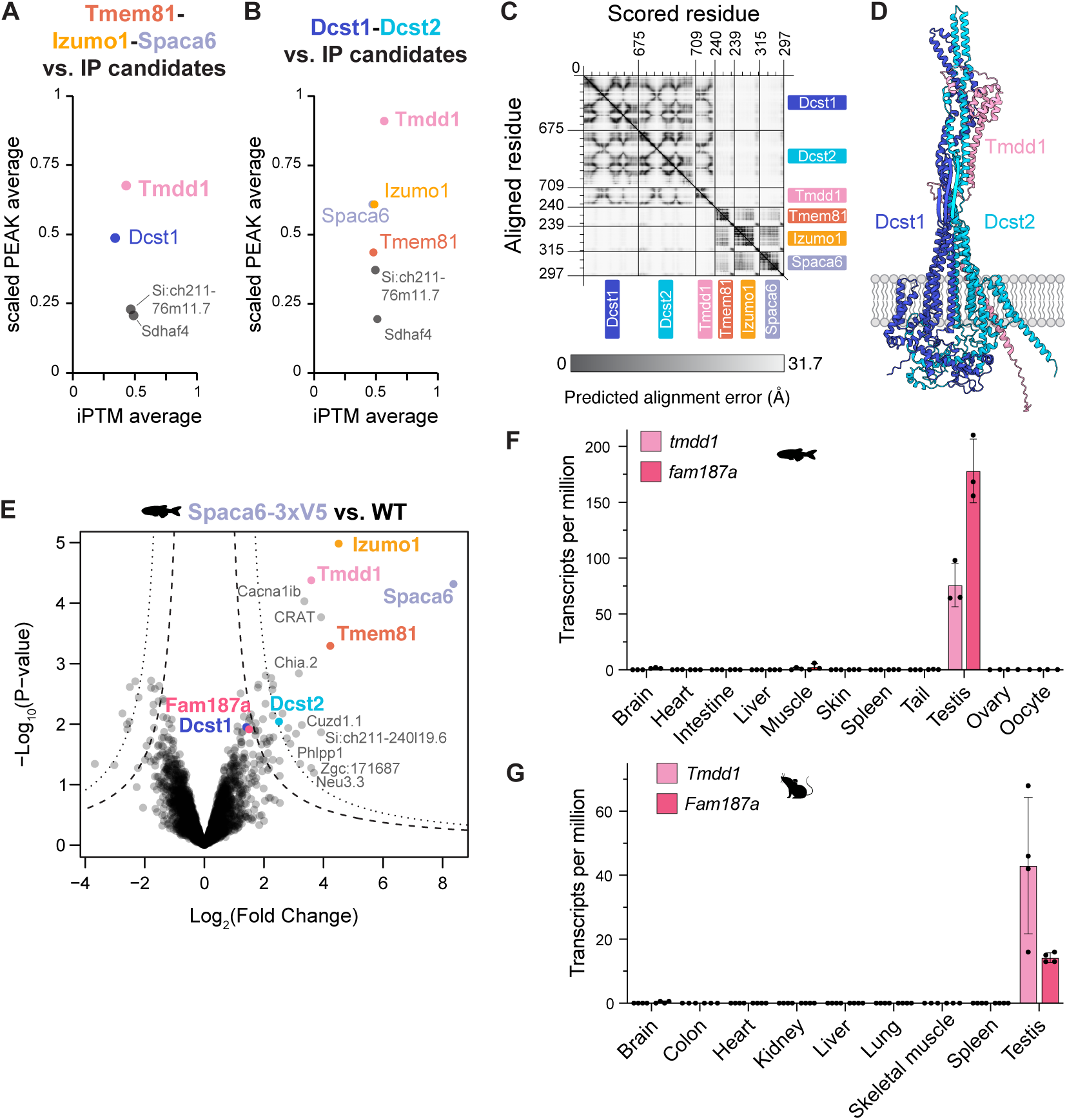
Evidence for SPARK complex formation in zebrafish and mouse. (**A, B**) AlphaFold screen predicting pairwise interactions between candidate factors significantly enriched in Tmem81-3xFLAG co-IP experiments from zebrafish sperm and Tmem81-Izumo1-Spaca6 (**A**) or Dcst1-Dcst2 (**B**). iPTM: interface predicted template modeling score. The scaled PEAK average score represents the minimal predicted alignment error (PAE) values in the interface between the candidate complexes and interactors on a scale from 0 (high PAE, low confidence) to 1 (low PAE, high confidence) (*34*). (C) PAE plot of the AlphaFold prediction of zebrafish Tmdd1 together with Dcst1/2 and Tmem81-Izumo1-Spaca6. Tmdd1 is confidently predicted to interact with Dcst1/2 but no interaction is predicted between Tmdd1-Dcst1-Dcst2 and Tmem81-Izumo1-Spaca6. (D) Predicted model of the zebrafish Dcst1-Dcst2-Tmdd1 trimer. iPTM: 0.64. (E) Volcano plots of differentially enriched proteins in V5 co-IP experiments from zebrafish sperm expressing Spaca6-3xV5 versus wild-type (WT) sperm. False discovery rate (FDR) thresholds are indicated as dotted (1% FDR) and dashed lines (5% FDR). Detected SPARK complex members are highlighted in color. (**F, G**) Expression of zebrafish *tmdd1* and *fam187a* mRNAs (**F**), and of mouse *Tmdd1* and *Fam187a* mRNAs (**G**) in adult tissues. RNA-seq data from: (*14*, *18*); mouse TPM values were retrieved from the Expression Atlas: https://www.ebi.ac.uk/gxa/home (*106*).

**Fig. S2.**
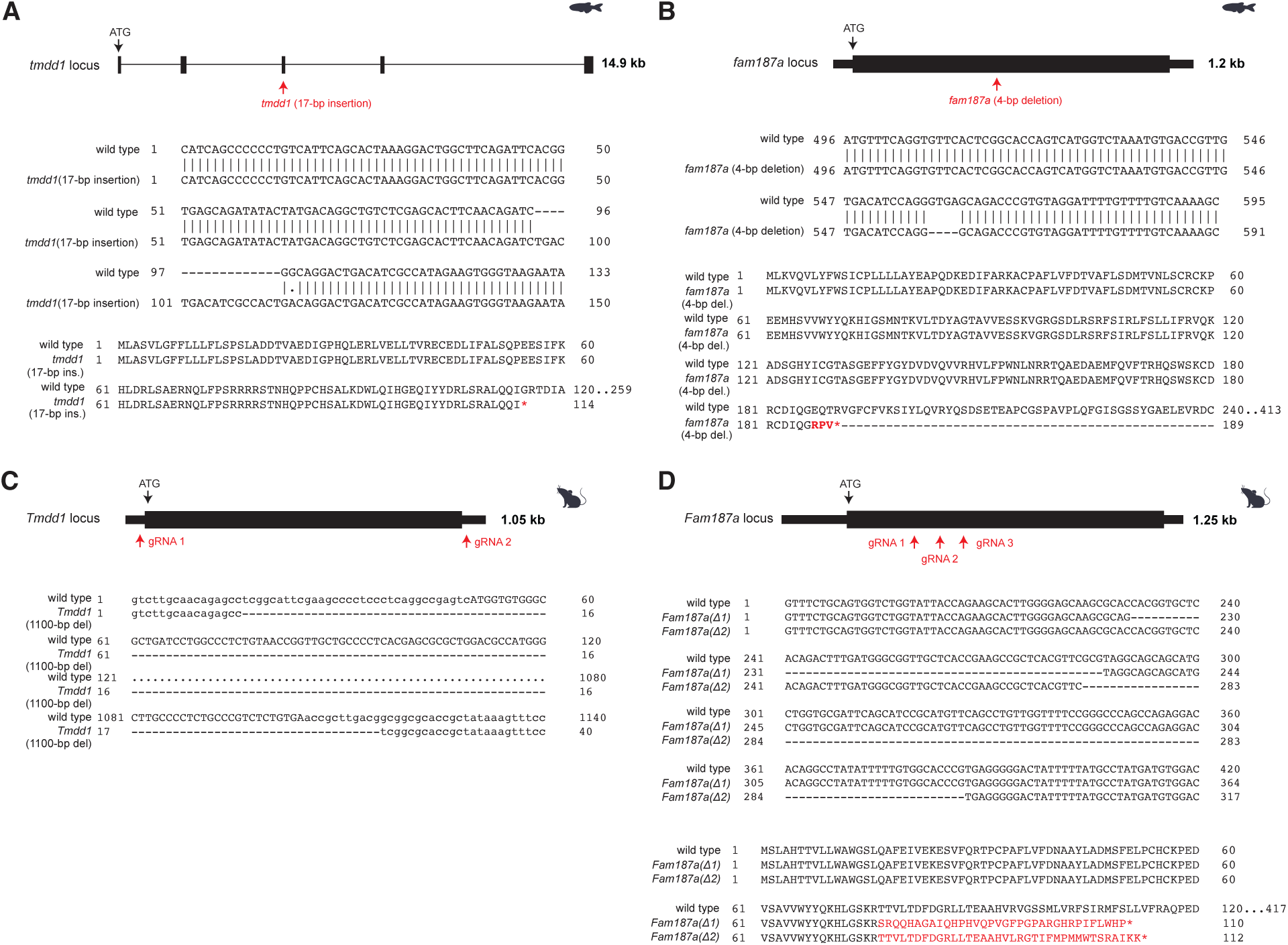
Establishment of *tmdd1* and *fam187a* mutants in zebrafish and mice. (A) Zebrafish *tmdd1* mutant. Top: scheme of the *tmdd1* locus in zebrafish indicating the location of the 17-bp insertion (boxes: exons; lines: introns). Middle: partial cDNA sequence alignment of exon 3 of wildtype and the *tmdd1* mutant. Bottom: Partial amino acid sequence alignment for wild-type and mutant Tmdd1. The 13-nt insertion in mutant *tmdd1* leads to a frameshift that results in a premature stop codon. (B) Zebrafish *fam187a* mutant. Top: scheme of the *fam187a* locus in zebrafish indicating the location of the 4-bp deletion (thick boxes: exons; thin boxes: UTRs). Middle: partial cDNA sequence alignment of wildtype and the *fam187a* mutant. Bottom: Partial amino acid sequence alignment for wild-type and mutant Fam187a. The 4-nt deletion in mutant *fam187a* leads to a frameshift that results in a premature stop codon. (C) Mouse *Tmdd1* mutant. Top: KO strategy of mouse *Tmdd1* indicating the location of 2 single-guide RNAs (sgRNAs). Bottom: Partial genomic sequence alignment of wildtype and the *Tmdd1* mutant, resulting in a 1100-bp deletion at the *Tmdd1* locus, which removes the entire *Tmdd1* coding sequence. (D) Mouse *Fam187a* mutant. Top: KO strategy of mouse *Fam187a* using 3 single-guide RNAs (sgRNAs). Middle: Partial genomic sequence alignment of wildtype and the *Fam187a* mutants (*Fam187a ^Δ1^* and *Fam187a ^Δ2^*). Bottom: Partial amino acid sequence alignment for wild-type and mutant FAM187a. Deletions in mutants *Fam187a ^Δ1^* and *Fam187a ^Δ2^* lead to frameshifts that result in premature stop codons.

**Fig. S3.**
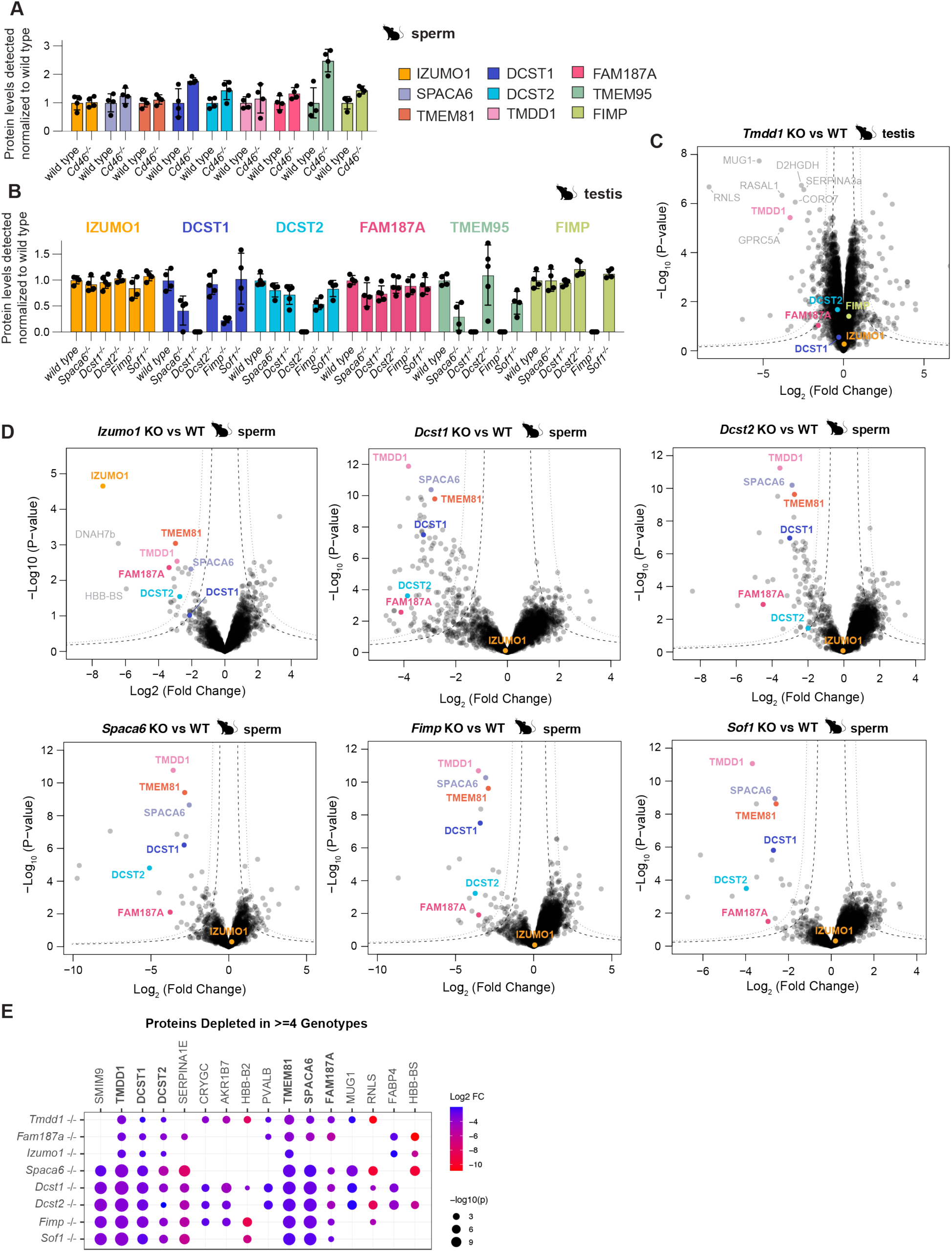
SPARK complex protein measurements in mouse sperm and testis. **(A)** PRM-MS of SPARK complex members of wild-type and *Cd46* KO mature mouse sperm (n = 4 / genotype) normalized to four constant proteins. SOF1 could not be detected by PRM–MS due to lack of suitable peptides. **(B)** PRM-MS of SPARK complex members of wild-type, *Spaca6* KO, *Dcst1* KO, *Dcst2* KO, *Tmdd1* KO, *Fimp* KO, and *Sof1* KO testis samples (n = 4 / genotype) normalized to four constant proteins. TMDD1, TMEM81, and SPACA6 were not confidently measured from testis samples and are therefore not included. SOF1 could not be detected by PRM–MS due to lack of suitable peptides. **(C)** Volcano plots of differentially regulated proteins in wild-type (WT) versus *Tmdd1* KO testis samples. Detected SPARK complex members are highlighted in color. False discovery rate (FDR) thresholds are indicated as dotted (1% FDR) and dashed lines (5% FDR). **(D)** Volcano plots of differentially regulated proteins in wild-type (WT) versus *Izumo1* KO, *Dcst1* KO, *Dcst2* KO, *Spaca6* KO, *Fimp* KO, and *Sof1* KO mature sperm samples. Detected SPARK complex members are highlighted in color. False discovery rate (FDR) thresholds are indicated as dotted (1% FDR) and dashed lines (5% FDR). **(E)** Bubble plot of significantly depleted proteins (Log2 Fold Change < -2, *p*-value < 0.05) in sperm samples of at least four distinct genotypes. Color gradient indicates the magnitude of depletion (Log2 Fold Change), bubble size corresponds to the statistical significance (-log10(*p*)). Proteins and genotypes were organized using hierarchical clustering (Euclidean distance, complete linkage) based on their Log2 Fold Change profiles. SPARK complex members are in bold.

**Fig. S4.**
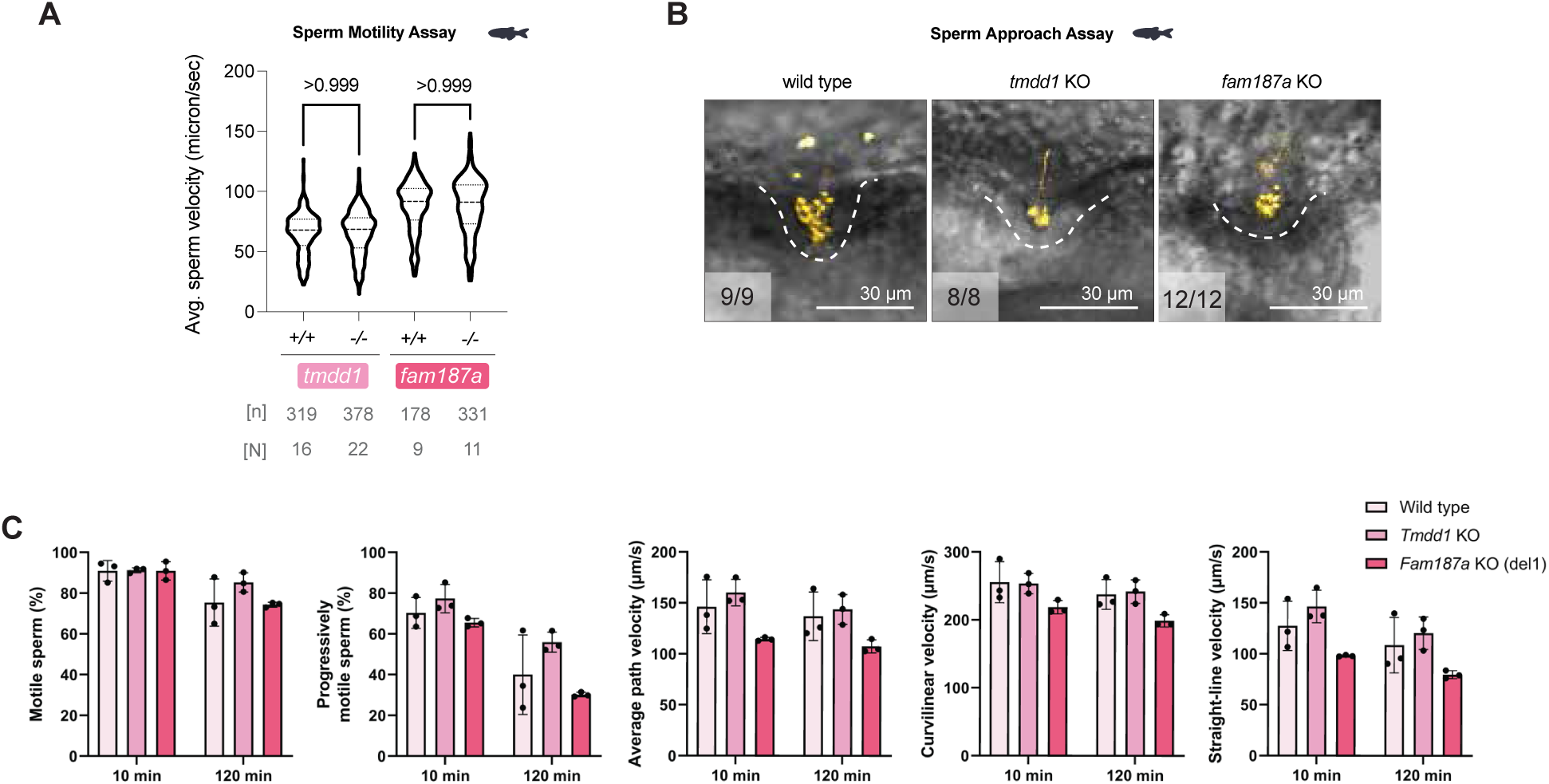
Predicted protein structures, conservation, and mutant phenotypic characterization. **(A)** Average sperm motility for zebrafish wild-type (+/+), *tmdd1* and *fam187a* KO (-/-) sperm. Violin plot indicates quartiles. *P-*values are calculated by Mann-Whitney test. **(B)** Zebrafish sperm approaching the micropyle of wild-type eggs. Representative images of MitoTracker-labeled wild-type (left), *tmdd1* KO (middle), and *fam187a* KO (right) zebrafish sperm (yellow) approaching the micropyle (white dashed lines). **(C)** Sperm motility, progressive motility, and swimming velocity in wild-type, *Tmdd1 -/-*, and *Fam187a ^Δ1^* -/- male mice.

**Fig. S5.**
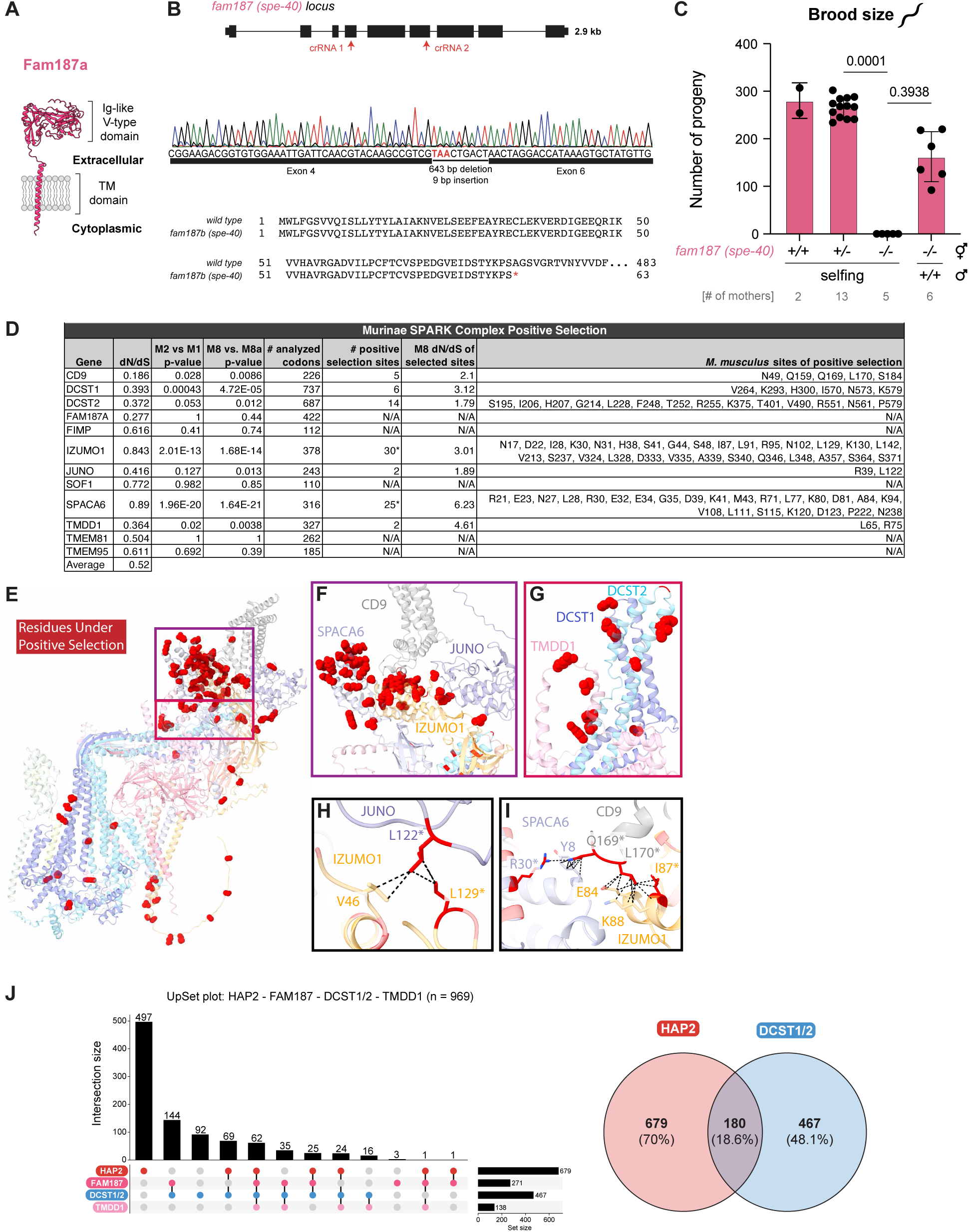
Evolutionary conservation of TMDD1 and FAM187A. **(A)** AlphaFold predicted structure and domains of zebrafish Fam187a. **(B)** *C. elegans fam187/spe-40* mutant. Top: genomic locus of the *fam187/spe-40* gene (Y37E11AR.7). The two crRNAs used for CRISPR/Cas9 editing are indicated. Middle: Genomic sequence trace of the *fam187/spe-40* mutant, revealing the introduction of a 634-bp deletion with a premature stop codon. Bottom: Amino acid sequence alignment of wild-type and mutant Fam187/Spe-40. The premature stop codon leads to termination after Ser63 in the mutant. **(C)** Brood size assay to assess the fertility of the *C. elegans fam187*/*spe-40* mutant. Number of self-progeny of hermaphrodites of the indicated genotypes (+/+, wildtype; +/-, heterozygous; -/-, KO), or number of progeny of homozygous mutant hermaphrodites mated with wild-type males. Bar graphs represent mean ± SD. *P*-values are calculated by Kruskal-Wallis test with Dunn’s multiple comparisons test. **(D)** Components of the SPARK complex evolve under positive (diversifying) selection in *Murinae*. All ten components of SPARK as well as the egg receptors CD9 and JUNO were tested for the presence of both gene-level and site-specific positive selection. High SPARK gene dN/dS values suggest the presence of adaptive pressure. **(E)** The predicted *M. musculus* SPARK complex ribbon structure bound to CD9 and JUNO with residues under positive selection identified as red atoms. (**F-G**) Zoom views from the whole SPARK complex show concentrated regions of positive selection at the IZUMO1/SPACA6 distal tip (**F**) as well as the DCST1/2 and TMDD1 tip (**G**). (**H-I**) Zoom views of physically interacting residues under positive selection between murine JUNO and IZUMO1 (**H**) and SPACA6, CD9, and IZUMO1 (**I**) suggest concerted evolution between egg and sperm proteins shapes the SPARK evolutionary trajectory. (**J**) Upset plot of HAP2, FAM187, DCST1/2, TMDD1 in invertebrates, showing the frequency of co-occurrence of the indicated proteins in invertebrates. A Venn-diagram (right) of species containing HAP2 (red) or DCST1/2 (blue) shows limited overlap between the two groups, revealing relatively few species (180; 18.6%) that have both HAP2 and DCST1/2.

**Fig. S6.**
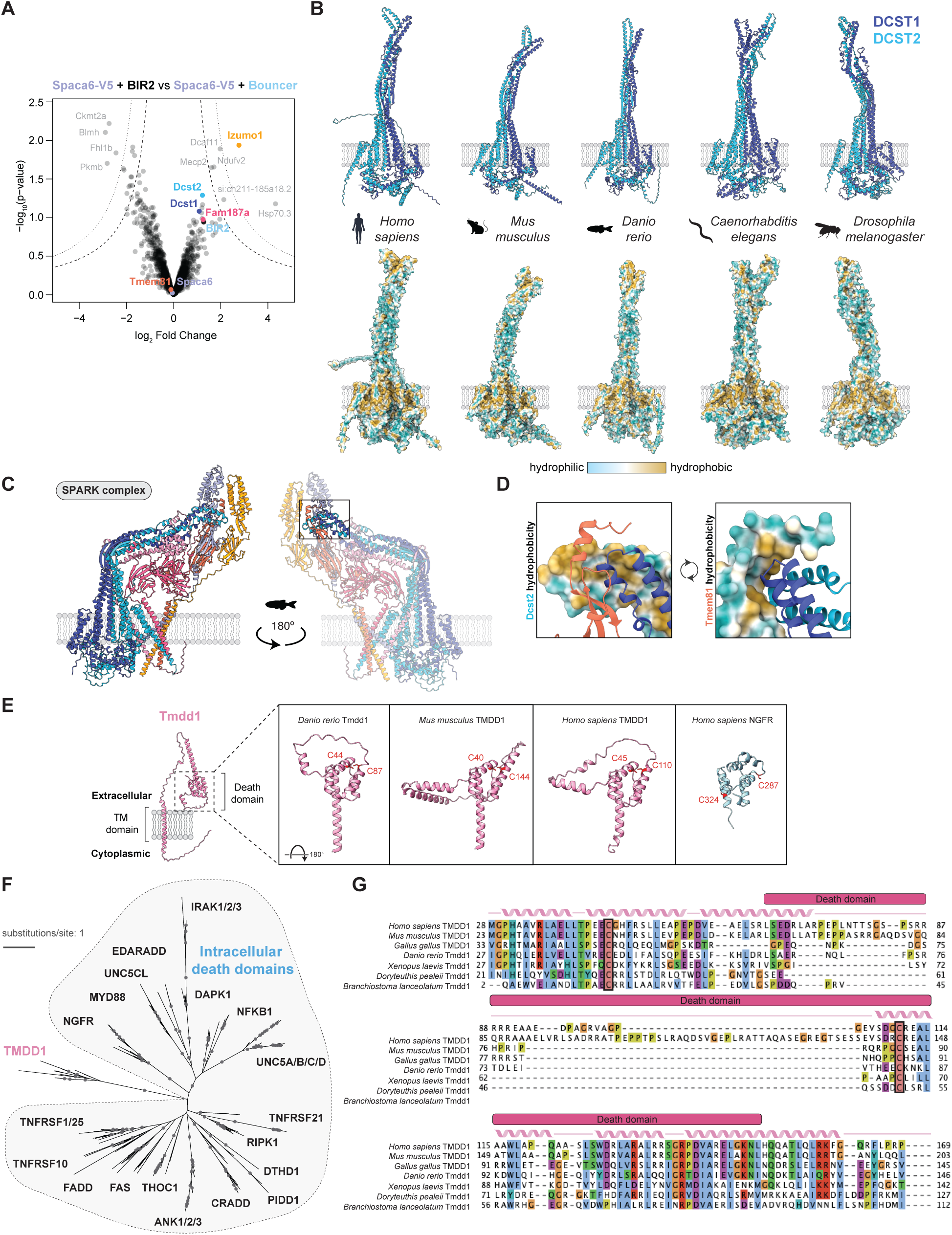
SPARK complex structural predictions suggest a conformation-driven activation mechanism. **A)** Volcano plots of differentially enriched proteins in V5-co-IP experiments from zebrafish sperm expressing Spaca6-3xV5, preincubated with either recombinant BIR2 (solubility tag) or BIR2-Bouncer. False discovery rate (FDR) thresholds are indicated as dotted (1% FDR) and dashed lines (5% FDR). The only significantly enriched SPARK complex member upon pre-incubation of sperm with Bouncer protein is Izumo1. **B)** AlphaFold models of the DCST1/2 dimer in different species. Top: ribbon diagram; bottom: surface representation colored by hydrophobicity. **C)** The N-terminal domain of Tmem81 is predicted to interact with the tip of Dcst1/2. Shown is the zebrafish SPARK complex. **D)** The hydrophobic tip of Dcst1/2 is predicted to interact with hydrophobic residues of Tmem81 (zoom-in of C). Top: surface representation of Dcst2 colored by hydrophobicity and ribbon model of Tmem81 and Dcst1; bottom: surface representation of Tmem81 colored by hydrophobicity and ribbon model of Dcst1 and Dcst2. **E)** Domain architecture of TMDD1. Left: AlphaFold predicted structure of zebrafish Tmdd1 (without signal peptide). Transmembrane (TM) domains, death domain and Ig-like V-type domains are labeled. Right: death domains of zebrafish, mouse and human TMDD1 compared with the death domain of human NGFR. TMDD1 has a unique disulfide bond (cysteine residues in red) that is not conserved in other death domain containing proteins, supporting its divergence from other death domain proteins. **F)** Unrooted phylogenetic tree of death domain containing proteins. Bootstrap values > 94 are indicated by filled gray circles. Branch lengths represent the inferred number of amino acid substitutions per site. The TMDD1 protein family stands out as the only family having an extracellular death domain, while all other death domain containing proteins have intracellular death domains (grey shading). **G)** Protein sequence alignment of TMDD1 with predicted secondary structure (ribbons: alpha helices; lines: unstructured) and death domain annotation. The conserved cysteines forming the disulfide bridge are highlighted. Amino acids reflect positions in full-length sequences.

